# Statistics and simulation of growth of single bacterial cells: illustrations with *B. subtilis* and *E. coli*

**DOI:** 10.1101/157545

**Authors:** Johan H. van Heerden, Hermannus Kempe, Anne Doerr, Timo Maarleveld, Niclas Nordholt, Frank J. Bruggeman

## Abstract

The inherent stochasticity of molecular reactions prevents us from predicting the exact state of single-cells in a population. However, when a population grows at steady-state, the probability to observe a cell with particular combinations of properties is fixed. Here we validate and exploit existing theory on the statistics of single-cell growth in order to predict the probability of phenotypic characteristics such as cell-cycle times, volumes, accuracy of division and cell-age distributions, using real-time imaging data for *Escherichia coli* and *Bacillus subtilis*. Our results show that single-cell growth-statistics can accurately be predicted from a few basic measurements. These equations relate different phenotypic characteristics, and can therefore be used in consistency tests of experimental single-cell growth data and prediction of single-cell statistics. We also exploit these statistical relations in the development of a fast stochastic-simulation algorithm of single-cell growth and protein expression. This algorithm greatly reduces computational burden, by recovering the statistics of growing cell-populations from the simulation of only one of its lineages. Our approach is validated by comparison of simulations and experimental data. This work illustrates a methodology for the prediction, analysis and tests of consistency of single-cell growth and protein expression data from a few basic statistical principles.

---

Thousands of biochemical reactions are required for bacterial growth and division. Some of them operate in a regime where they are susceptible to stochastic fluctuations in the concentrations of their reactants and regulators^1,2^. These fluctuations can be amplified by molecular networks^3^, in particular by those with positive feedback circuitry and propagated to the cellular level, causing variations in the growth characteristics^4^ (birth size, division size, generation time etc.) and molecular content^5^ of individual cells. A reverberating coupling exists between the molecular composition of a cell and its growth behaviour^4^, where fluctuations at a cellular level can in turn cause cell-to-cell variations at the level of molecules and reaction activities. For instance, uneven cell division causes size differences between cells such that their protein content and reaction rates vary^4,6,7^. The fluctuating copy number of a particular molecule in a cell, over a time period of several bacterial cell cycles, is therefore the outcome of (stochastic) biochemical and cell-growth processes^8–10^.

Coupled molecular and cellular fluctuations are associated with many surprising phenomena in single-cell biology^10–14^. This complex feedback circuitry generates cell-to-cell variability in a population of isogenic cells, which may result in individual cells transiting to different phenotypic states when conditions change^11–13,15,16^. Examples include adaptive phenotypic-diversification of populations of cells, e.g. the emergence of antibiotics-tolerant persister cells^2^ and bacterial-cell differentiation^16^. Acquiring a predictive understanding of these phenomena is one of the current challenges in single-cell physiology^17^, with direct applications in biotechnology^18^ and medical microbiology^19,20^. Disentangling causes and effects, using a stochastic framework, is a major challenge for single-cell physiology^21,22^ and methods need to be developed that can quantify the contributions made by the stochastic biochemistry of molecular circuits and cellular growth, including theory (e.g. variance decomposition) and simulation^6,21–23^.

In contrast to the complexity of molecular and cellular processes at a single-cell level, the macroscopic, population-level properties of bacterial cultures are much easier to quantify and predict. In fact, the properties of bacterial cultures at balanced growth follow surprisingly simple ‘laws’. Examples are the relations of Malloe-Schaechter-Kjeldgaard^24^, Monod^25^ and Pirt^26^, which were developed in the 1950-70s. In that same period, a statistical theory was derived for the behaviour of single cells at balanced growth – a ‘microscopic’ growth theory ^27–29^. It describes statistical relations between growth properties of single cells, such as birth and division volumes, generation times, growth rates, and daughter-mother volume ratios. With this theory, quantitative descriptions of populations can account for inter-individual variations in the physiological parameters (i.e. their distributions) of asynchronously growing cells. In the current work, we validate this statistical theory with real-time imaging of bacterial growth and fluorescent-protein expression, using time-lapse fluorescence microscopy. We show that relations between different phenotypic characteristics can be used in consistency tests of experimental data of single-cell growth, and the prediction of single-cell statistics. We then exploit these robust correlations to develop a fast and predictive stochastic simulation algorithm of single-cell growth and protein expression. Together, the statistical relations and the stochastic simulation algorithm offer a methodology for prediction, interpretation and tests of consistency of experimental data of the stochasticity of single-cell growth and molecular circuit dynamics.

## Results

### Describing the growth characteristics of single cells at balanced growth

In the following sections, we will validate the statistical relations captured by microscopic growth theory^27^’^29–31^ and show how this framework allows for a comprehensive quantitative description of single-cell growth characteristics from a limited set of physiological single-cell growth-measures. The relations we will validate make use of several concepts, which we will first explain.

For any single cell, its age can be defined as the time elapsed since its birth. At birth, the ‘cell age’ is zero and all cellular properties are birth properties, e.g. birth-volumes, -lengths, -widths. Every cell has as its maximal age its ‘generation time’ (the generation time is also sometimes referred to as the interdivision or cell-cycle time), which is the time at which it divides and has attained its ‘division size’. The specific rate with which this volume increases, *d* ln*V/dt* is called the ‘instantaneous specific growth rate’. When the division volume is reached, a cell divides and partitions its volume and molecular content to yield two new cells. The dividing cell is referred to as the mother cell and the two newly formed cells as the daughters, which are sisters. The size and molecular content of each of the two daughters can be expressed as a fraction of their mother’s to capture cell-division variation. All these properties together (cell age, generation time, birth and division sizes, daughter-to-mother ratio’s and instantaneous growth rate), capture the essential information needed for a microscopic theory of growth^27,29–31^ (see Fig. 1 and the appendix for additional information related to concepts of single cell growth).

**Figure 1.**
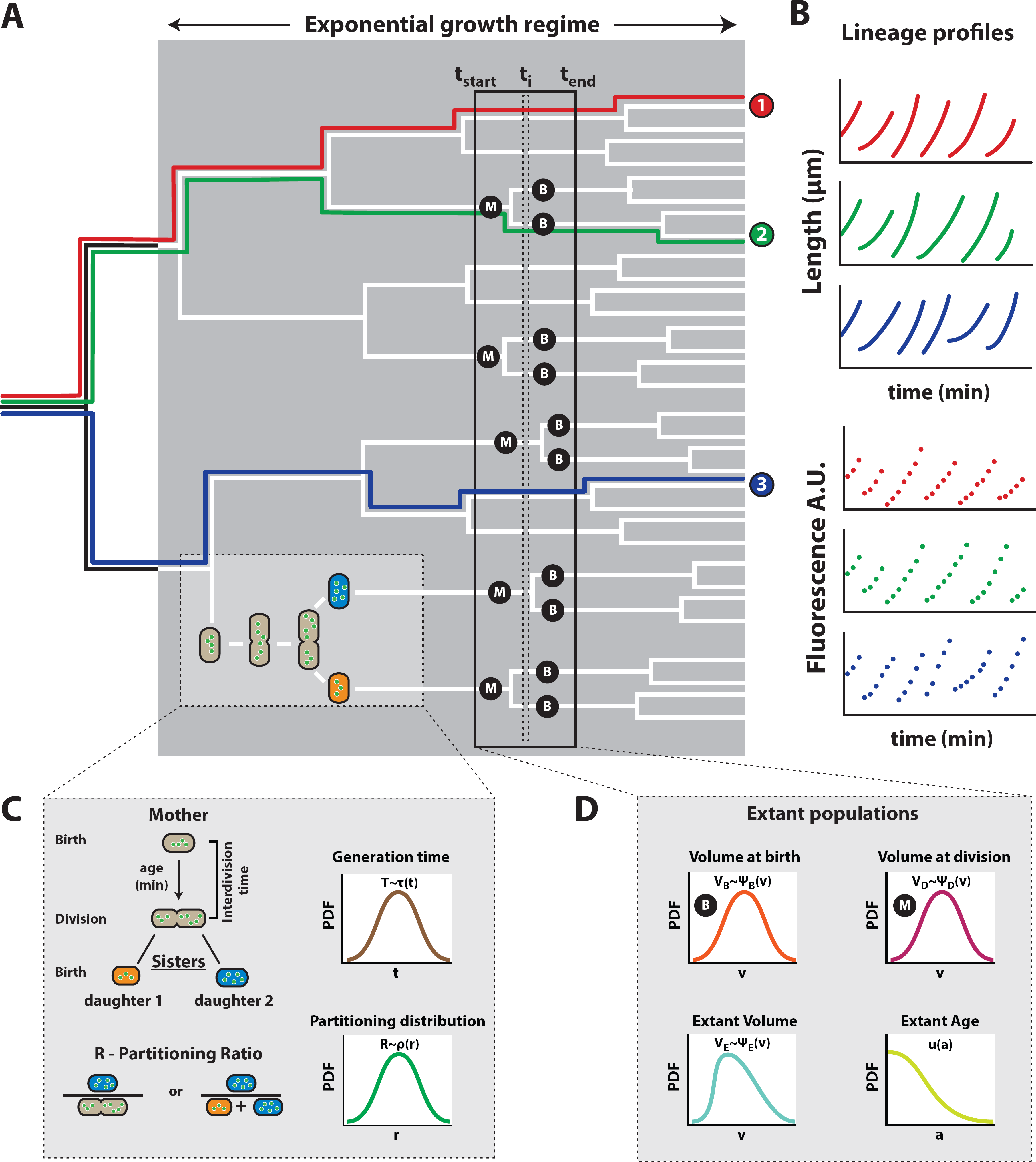
Growth characteristics and concepts of single cells in a population at balanced growth. (A) The formation of a microcolony from a single ancestral cell can be represented as a lineage tree. In such a tree, time runs from left to right, horizontal lines represent the life lines of single cells, their total length equals the generation time of a cell, and vertical lines indicate cell divisions. (B) A lineage corresponds to the growth and division of single cells, that are all daughters from a specific ancestral cell. At specific time points along a lineage, the cell length and fluorescence can be measured. (C) After a cell-cycle duration, corresponding to the generation time of a (mother) cell, two daughter cells arise via imperfect cell division, giving rise to a probability to observe daughter cells that have obtained a certain fraction of their mother cell’s volume and molecular content. (D) At one given moment in time all extant cells have particular properties that follow probability distributions such as their birth volume, division volume, current volume and current age. Extant populations consist of cells that divide (mothers, M) and cells that are born (Babies, B).

In any population these single-cell growth-measures will show variation from cell to cell, necessitating a statistical framework to understand how population-level characteristics relate to single-cell phenotypes, and how different single-cell growth-measures depend on each other. Answers to these types of questions are greatly simplified when populations are studied under conditions of balanced growth. A defining characteristic of balanced growth is that the probabilities to observe cells with particular growth properties – their phenotype – are fixed and the associated probability distributions are therefore time invariant (See also Fig. S3). Importantly, the validity of the statistical relations captured by the microscopic growth theory rests strongly on the assumption that the population being described is at balanced growth.

Balanced growth, being a stationary process, has as a requirement that the specific growth rate of the population remains fixed over a time period that is several times longer than the mean generation time. As such, the single cell growth data we use to validate the microscopic growth theory^27,29-31^ was confirmed to meet this requirement. By individually tracking the growth of *B. subtilis* and *E. coli* cells on agar pads, we quantified the specific growth rate of the population from the increase in the total cell length of all monitored cells, and selected data from the time-window during which the growth rate remains fixed. We confirmed that the balanced-growth period lasted for several generations and that the probability distributions of growth measures are constant during in this window^27^ (see Fig. S3). All growth measurements of B. *subtilis* can be found in Fig. 2 (discussed below) and those of *E. coli* are shown in the Supplemental Information (Fig. S4).

**Figure 2.**
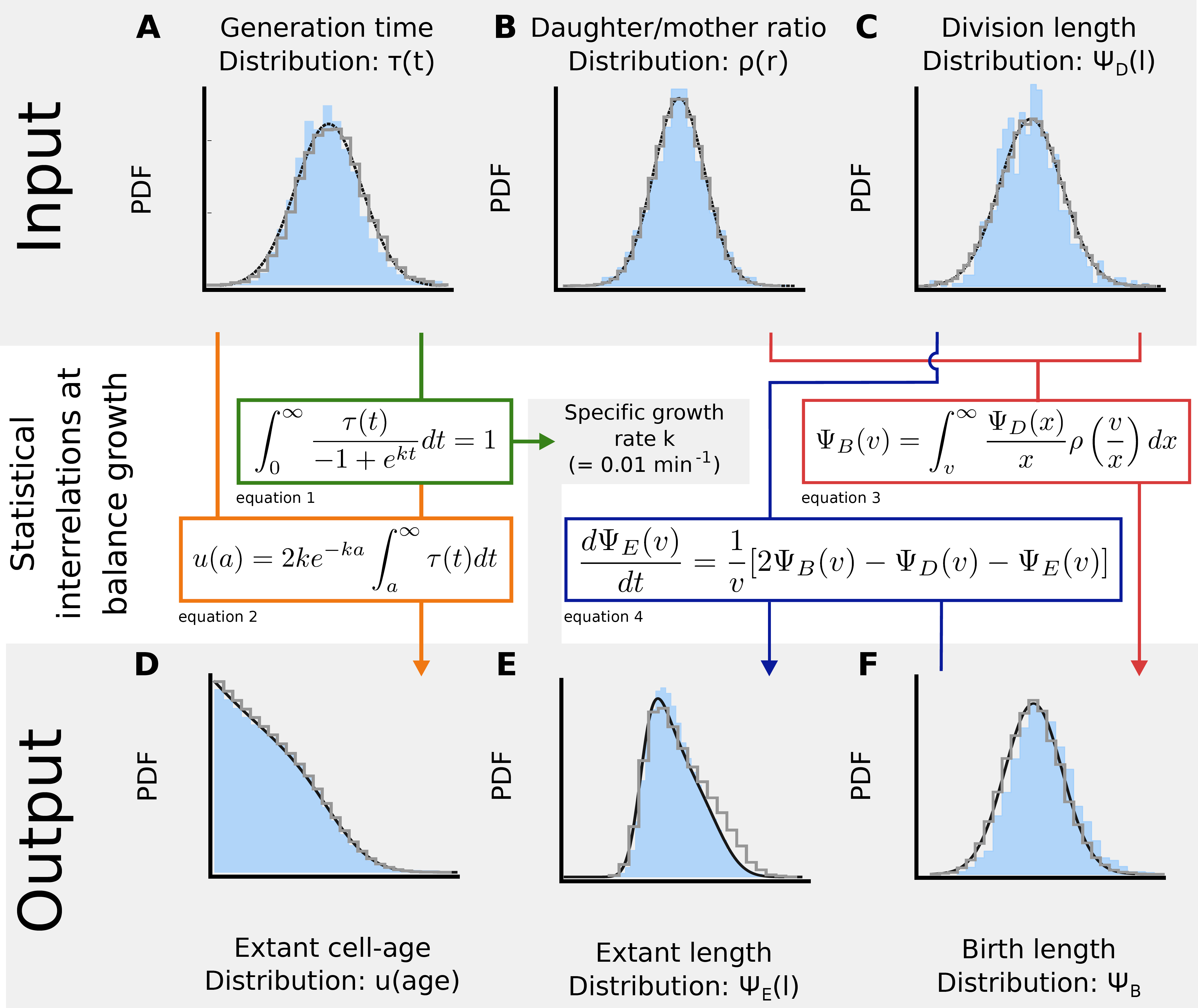
Validation of relations between growth characteristics at balanced growth for *B. subtilis.* Shown are results comparing the microscopic growth theory relations derived by Collins & Richmond^29^, Powell^28^ and Painter & Marr^27^ and experimental single-cell growth data. In this figure we validate those relations. The probability distributions obtained from experimental data are shown in blue, the validated theoretical relations are shown in the coloured boxes (eqs. 1 – 4), the predicted distributions are shown in black in (D), (E) and (F), and the results of our simulation algorithm (discussed in the final result section) are shown as grey histograms. In (A)-(C), black lines indicate fits. We calculated the population specific growth rate from the distribution of the generation times (A). The distribution of the cell ages (D; age is the time elapsed since birth) can be obtained from the generation time distribution (A) and the calculated specific growth rate of the population. The distribution of cell length of all cells that exist at a particular moment in time, the extant cells (E), can be obtained from the distribution of the birth lengths (F) and the division lengths (C). The distribution of birth lengths (F) can be obtained from the distribution of daughter-mother-volume ratios (B) and distribution of division lengths (C). Sample sizes: The sample sizes for the experimental data are 3637 extant cells, 2726 cells at birth and 1466 cells at division. Data for *E. coli* can be found in Fig. S4.

### The population growth rate calculated from single-cell generation times

The first statistical relation we validated allows for the calculation of the population growth rate (*k*) from the distribution of the generation times (Fig. 2, equation 1)^27,30,31^. In the macroscopic theory of bacterial growth of cell populations, the specific growth rate equals *k* = ln2/τ_g_, with τ_g_ as the generation time (also called the doubling time). Due to inter-individual variations in generation times, the macroscopic relation is inexact and the relation 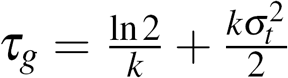 has been proposed as an improved approximation, with 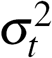 as the variance of the distribution of generation times^27^. The equation we used (derived in^27^; equation 1 in Fig. 2) obtains the exact value of the growth rate from the distribution of generation times.

We calculate from Painter & Marr’s relation a growth rate of 0.61 *hr^-1^.* When we compare this growth rate to the one we measured from the length increase of the population data (Fig. S1A), we find a difference of only 2.7%. From the measured growth rate, we can calculate the generation time, from ln2/*k*, which gives 70 *min.* If we use the approximate relation to calculate the generation time from the growth rate (0.01 *min*^−1^) and the measured generation-time variance of 388 *min*^2^, we find a generation time of 71 *min,* indicating that the approximate relation shows an error of 1.4%. We conclude that the measuredprobability distribution of the generation times indeed allows for a calculated value of the cell-population’s growth rate that is in agreement with an independent direct observation of the growth rate from the increase in length of the entire population of cells.

The generation-time distribution, τ(t), that we measured (Fig. 2A) can be accurately approximated by a normal distribution, which has also been observed by others^5,32-35^. The coefficient of variation of the generation time is 27%, which indicates that 16% of the cells have a generation time below 53 *min* and the generation time of the same percentage of cells exceeds 92 *min.*Significant variations in generation times therefore occur in steady-state growing bacterial cell populations. Other work has recently shown that the growth rate, the length increase during a cell cycle and the generation time are interrelated^5,32,36^.

### Obtaining the distribution of cell ages from the generation times of single cells

Since all cells grow asynchronously during balanced growth and because balanced growth is a stationary process, the probability that a cell is observed with a particular age – the time elapsed since its birth – is constant. The age of a cell ranges from 0 to its generation time, when it divides.

Painter & Marr^27^ showed that the distribution of the probabilities for the occurrence of a particular cell age can be obtained from the distribution of the generation times and the growth rate (Fig. 2, equation 2). Here we validate this theory by showing that a theoretically calculated age distribution, using Painter & Marr’s relation, closely matches the independently-observed cell age distribution (Fig. 2D), indicating that the theory holds for our dataset. This finding also confirms that, during the time window of data sampling, the cells were indeed growing balanced.

We observed that the mean age of extant *B. subtilis* cells equals 32 *min* (Fig. 2D). Since the mean generation time (the mean division age) equals 75 *min,* the average cell has a cell cycle progression of 43%. The cell-age distribution also indicates that a cell population in balanced growth contains more young than old cells, which is expected since a mother cell always divides into two daughter cells.

### The distribution of daughter-over-mother cell sizes calculated from the distributions of birth and division sizes

Powell^28^ derived a relation between the distributions of birth and division volumes and the distribution of daughter-over-mother volume ratios (Fig. 2, equation 3). This theory is in terms of the volume distributions of cells. Here we apply it to analyse cell length data (Fig. 2C). Throughout this paper, we treat length changes proportional to volume changes because the width of B. *subtilis* and *E. coli* cells remains roughly constant during growth (Fig. S2 and^5,37^). Therefore, the growth in cell volume is proportional to the growth in cell length.

We approximate the daughter-over-mother length ratio in our experimental data using the ratio of one daughter cell over the sum of her length and that of her sister (Fig. 2B). Since we monitor growth at 1-minute intervals, during which some growth takes place, a small discrepancy occurs between our estimate of the daughter-vs-mother length ratio and the real value (see also supplement information section 4).

For symmetrically dividing bacteria, such as *B. subtilis* and *E.coli,* the average daughter-over-mother length ratio is expected to be 0.5, which is reflected by our data (Fig. 2B). Application of the Powell relation (equation 3, Fig. 2) leads to a prediction of the birth length distribution, that closely matches the experimental data (Fig. 2F).

The distributions of division length, birth length and daughter-over-mother length ratio’s that we measured can all be accurately approximated by normal distributions. The daughter-over-mother length ratio has the expected mean of 0.5 and a coefficient of variation of 7%, which captures division noise. The variation in division and birth length is about twofold greater. The birth and division lengths of cells also correlate with each other (Fig. S7 and S6), which is in agreement with an adder-like size-homoeostasis mechanism described for both *B. subtilis* and *E. coli*^5^. Our data, however, deviates slightly from perfect adder behaviour, which could be due to the use of agar pads, where the formation of micro-colonies could lead to differences between cells in the centre and those at the periphery of the colony.

### Calculation of the length distribution of cells

Collins & Richmond^29^ published a relation between the growth rate as a function of cell volume and the probability distributions of daughter, mother and extant cell-volumes. When the cellular growth rate is fixed as a function of the cell volume, as it is for our data (see Fig. S9), the Collins-Richmond relation simplifies to equation 4 of Fig. 2. We used this relation to predict the extant cell length distribution from the length distributions at birth and division. Figure 2E indicates that the predicted and measured extant cell length distributions are in close agreement. Variability in the extant cell length distribution is due to the fact that both daughter and mother cells occur in the extant cell population, and their lengths (or volumes) vary on average by a factor of two. Any additional spread in this distribution arises from noise in distributions of the birth and division length.

### Exploiting the microscopic growth theory for inference of probability distributions from data

Now that we have concluded that the microscopic growth theory is in agreement with single cell growth data, we can consider its applications. One application is its use in consistency checks of experimental data, as we did in Fig. 2; in order to test whether measurements of different growth characteristics relate to each other as we theoretically expect them to. Another application of the microscopic growth theory is its use in the inference of distributions of growth characteristics of single cells when not all of them can be directly measured.

Consider, for instance, a growth experiment carried out in a shake flask, where the specific growth rate and the probability distribution of the (extant) cell volumes, shown in Fig. 3A was measured (there many non-microscopy based methods for measuring cell size distributions, e.g. the Coulter counter). We shall assume that the birth and division length distributions each follow a normal distribution with means that they differ by a factor two; an assumption that is in fact in agreement with our data (Fig. 2). This leads to three unknown parameters: two standard deviations and one mean. Those we can obtain by fitting the division and birth-length distributions to the measured extant length distribution, using the Collins & Richmond relation (equation 4 in Fig. 2), parameterised with the measured growth rate. In this way, the probability distributions for the birth and division volume can be inferred, those are the solid lines in Fig. 3B and C.

**Figure 3.**
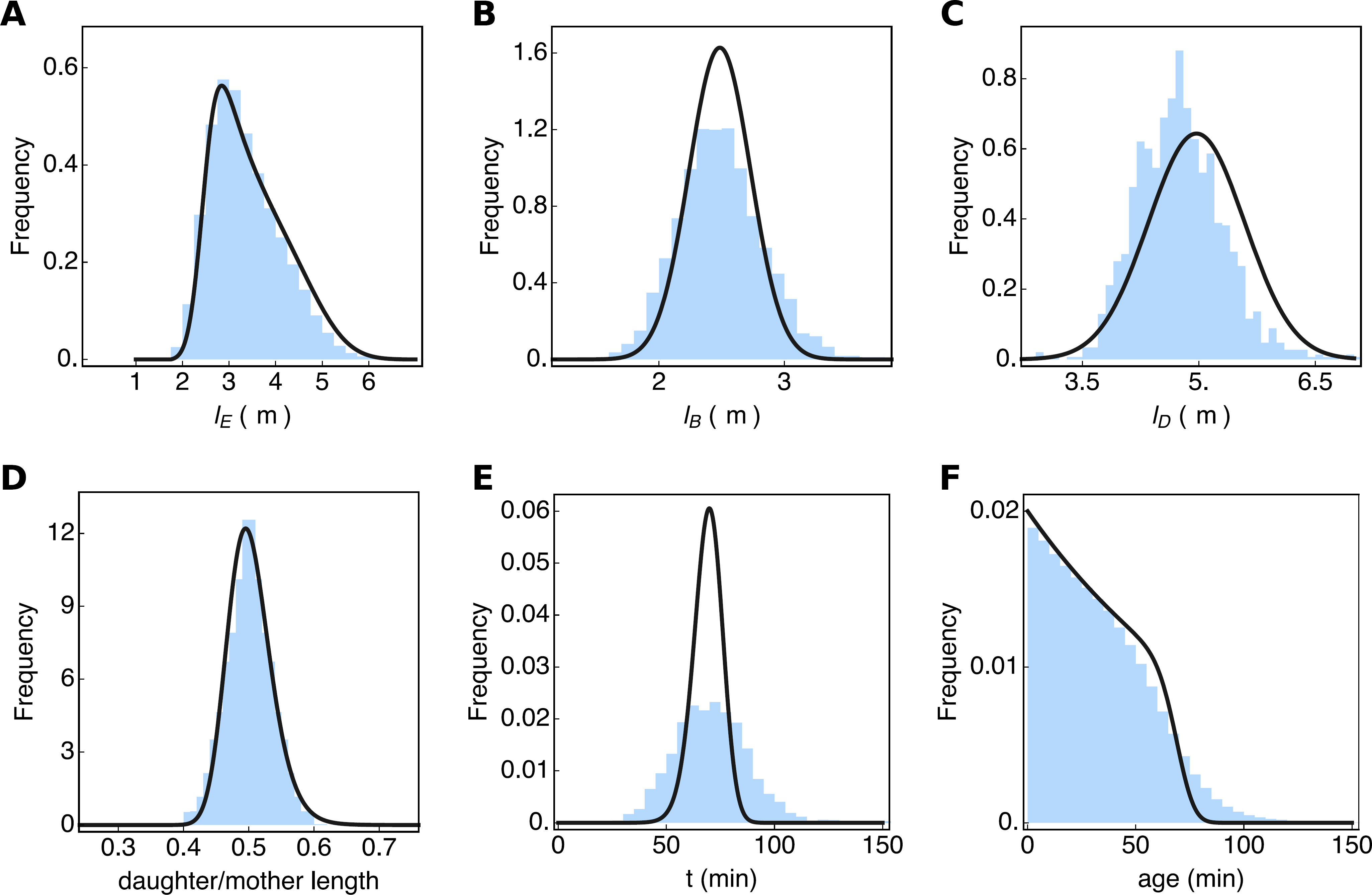
Inference of probability distributions of single-cell growth characteristics from experimental data. In this figure, the blue histograms are the measured data (also shown in Fig. 2) and the black lines are inferred from the data. (A) Shows the extant cell-length distribution. The cell length distributions at birth (B) and division (C) were obtained by a fit of the Richmond & Collins relation (equation 4 of Fig. 2) to the extant cell length distribution, given the measured growth rate. The distribution of the ratio of the daughter cell length over the mother cell length (D). It was obtained from the distributions shown in (B) and (C), assuming a correlation between cell length at birth and division of 0.85. By assuming deterministic exponential growth, the distribution of generation times (E), was predicted from the length ratio distribution (D). The distribution of cell ages (F) was calculated from the generation time distribution (E), using equation 2 from Fig. 2. Only the predicted generation time distribution shows a significant discrepancy with the experimental data, which we discuss in the main text.

To determine the distribution of the daughter over mother lengths, p *(r)* with *r = l_b_/l_d_*, we use a mathematical relation that expresses this distribution in terms of the joint distribution of *l*_*b*_ and *l*_*d*_ values^38^. We assume this joint distribution to be a bivariate normal distribution, which is in agreement with the data shown in Fig. 3B and C. We still need to assume a correlation coefficient for *l*_*d*_ and *l*_*b*_ values, which we can do by assuming a sizer, adder or timer model of cell growth^5,39^. When we assume an adder then the correlation coefficient will be close to 1. If we take a correlation coefficient of 0.88, we obtain a perfect fit of the estimated distribution of daughter over mother lengths with the experimental data (Fig. 3D), indicating this estimation method works. The fit quality is, however, strongly dependent on the exact value of the correlation coefficient between *l*_*d*_ and *l_b_*.

At balanced growth, with a fixed growth rate along the cell cycle (assumption 3), the following relation holds 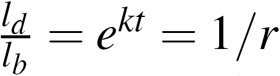 with *k* as the growth rate and *t* as the generation time. We can determine the distribution of generation times, t, from the distribution of the daughter-over-mother volumes, using the ‘change of random variable method’, which leads to the relation τ*(t) = ke^-kt^ρ (e^−kt^)*. This allows us to determine the generation time distribution from the distribution of daughter-over-mother lengths. We show the result in Fig. 3E. Clearly, the fit quality is poor, which is due to the fact that the assumed deterministic growth model *l_d_ = l_b_e^kt^* misses a stochastic component. What this component is, is unclear at the moment, but it is very likely that *k* depends *l*, such that the growth rates of smaller and larger newborn cells differ at birth.

Finally, the cell-age distribution can be determined from the estimated generation time τ*(t)* distribution (Fig. 3E), using equation 2 of Fig. 2. The resulting correspondence of the estimated distribution and the experimental data is shown in Fig. 3F. The fit is not perfect, due to the poor estimation of the generation distribution (Fig. 3E), but it is still acceptable given that it is derived from very basic assumptions and a few measurements.

### Exploiting the microscopic growth theory in a stochastic simulation algorithm

Another interesting aspect of the microscopic growth theory is its potential utility in the development of a fast stochastic simulation-algorithm of single-cell growth and molecular circuit stochasticity at balanced growth. Stochastic simulations of single-cell growth are very challenging, because of a large computational burden; they have to simulate tens of thousands of cells as they progress through their cell cycle and divide (giving rise to the expanding lineage tree). To circumvent this computational bottleneck, many existing approaches simulate only a single lineage, but do not retrieve the growth statistics of the entire lineage tree, thereby limiting meaningful comparisons between simulations and experimental data.

The algorithm that we developed simulates growth and molecular stochasticity of a single specific lineage, from which the statistics of the entire lineage tree can be calculated using the microscopic growth theory. First, we will briefly outline this algorithm (details are provided in the Supplemental Information, Section 7). Following this, we validate the algorithm using the experimental data introduced above, including single-cell data of a fluorescent gene expression reporter.

The input of the simulation algorithm for single-cell growth is based on several of the distributions of growth measures, quantified (or assumed) at balanced growth (Fig. 2). We start by calculating an initial cell volume, a birth volume *v*(0), from the product of two random variables, one sampled from the division-volume distribution, Ψ*_D_(v)* (Fig. 2C), and another from the division-ratio distribution, ρ *(r)* (Fig. 2B). Next, given our data, we assume exponential growth for single cells (Fig. S1), with a constant, specific growth rate that is equal to the population growth rate (k in Fig. 2)). From the difference between the division and birth volume, and the specific growth rate *k*, we can calculate the generation time (using 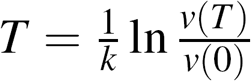). At the generation time, *T*, the cell divides into two daughter cells. One daughter receives a volume *v*_*d*1_ (0) = *rv(T)* (the value of *r* is drawn from the division-ratio distribution), which means that the second daughter’s volume will equal *v*_*d*2_(0) = *v(T) – v_d1_* (0).

Since our algorithm only simulates a single lineage, and not the whole lineage tree, we track only one of the two daughter cells, which makes the algorithm very computationally efficient. The current limitation of the algorithm is that it cannot deal with proteins that set the growth rate. Therefore, it simulates the stochasticity of biochemical circuits, while they are embedded in a cell that is growing in a stochastic manner, in line with the growth-statistics relations of balanced growth.

The simulation algorithm simulates growth of cells that grow according to sizer and ‘sizer-like adder’ mechanisms, which is how *B. subtilis* and *E. coli* grow (Fig. S6 and S7). The algorithm can also be used to simulate cells that grow as pure adders^5^. For adders, the volume at division is determined by the addition of a birth volume (drawn from the *V_B_* distribution) and an added volume value (drawn from a Δ*V* distribution). When it is assumed that the mother cell divides perfectly to yield two equally sized daughter cells, the algorithm simulates cell growth of a perfect sizer. The ratio between the partitioning variability and the division volume variability determines the slope in the (Δ*l|l_b_*)-vs-*l_b_* plot, and for the algorithm it falls between ‑0.2 and ‑1 (in accordance with data Fig. S6).

To be able to derive the statistics of the entire lineage tree from a single simulated lineage, a specific daughter cell is selected. The daughter that is selected is chosen with a probability that equals the fraction of descendants it is expected to contribute to the population (see Supplemental information section 7.1). After daughter-cell selection, we start simulating the next cell cycle, by sampling a new birth volume, etc. We repeat these steps until the lineage simulation has generated data for stationary distributions of growth measures. The statistics of the entire lineage tree can be obtained by exploiting the cell-age distribution as explained in the Supplemental Information.

We implemented this algorithm in StOChPy^23^, which is available for download from http://stochpy.sf.net. A full description of the cell growth algorithm and how it is coupled with a stochastic simulation algorithm for biochemical networks can be found in the Supplemental Information (Section 7). Importantly, we show that the stochastic simulation algorithm is in agreement with analytically solvable models (Supplemental Information). In the next section, we compare simulation results with experimental data.

### Validating the simulation algorithm with single-cell growth and protein expression data of *B. subtilis* and *E.coli*

The *B. subtilis* and *E. coli* strains that we used in our time-lapse microscopy experiments (Fig. 2 and Fig. S4) each expressed a genome-integrated, constitutively-expressed fluorescent protein. We used the fluorescence values of single cells as a read out of single-cell protein expression. We tested whether the stochastic simulation algorithm could reproduce single-cell growth and expression data (Fig. 4).

**Figure 4.**
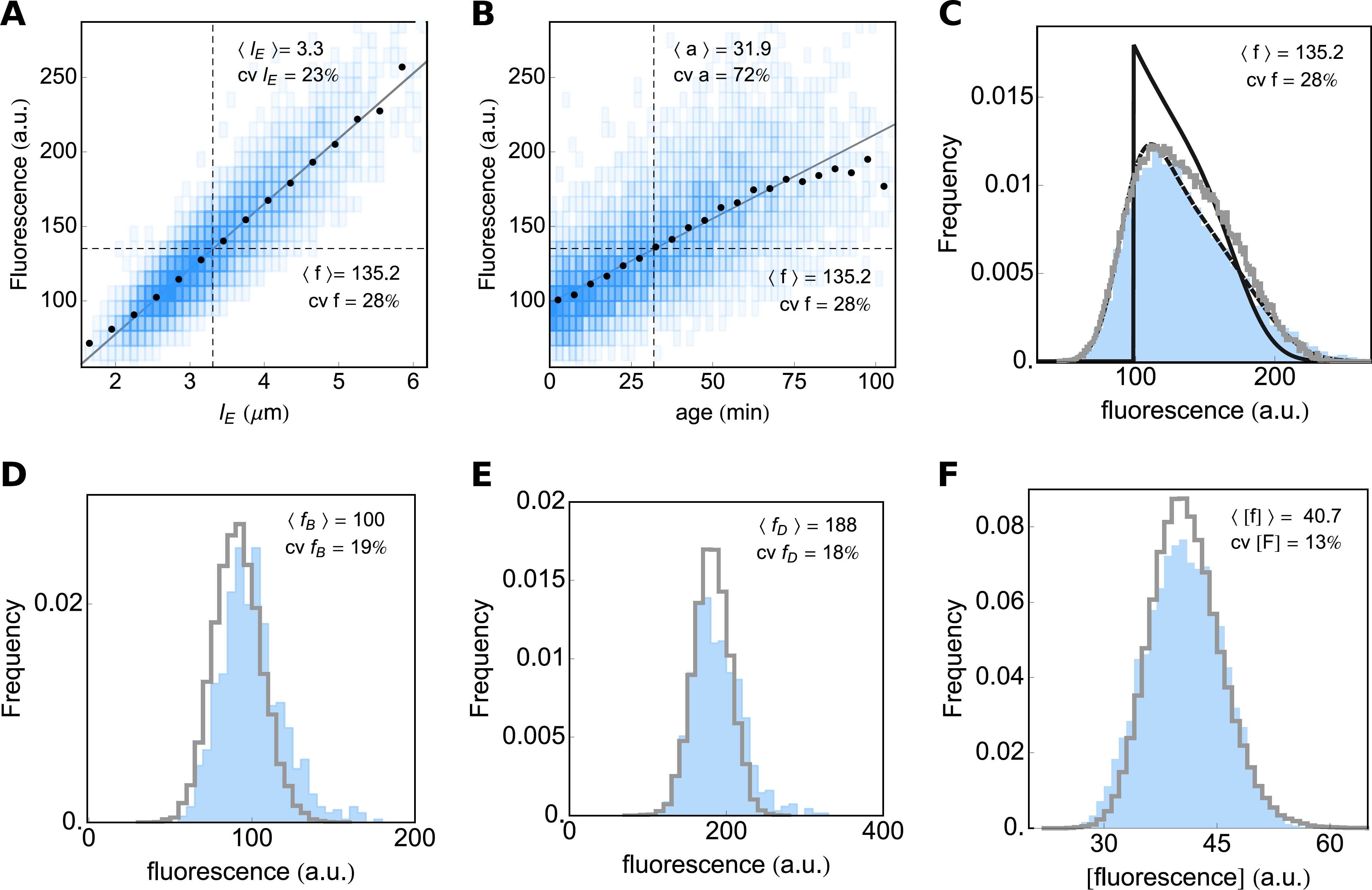
Expression data of the reporter gene of B. *subtilis.* Fluorescent protein expression scales linearly with cell length (A) and cell age (B), but the correlation is weaker for age. By transforming the extant length (Fig. 2E) and age (Fig. 2D) distributions with the linear relation between length and fluorescence and between age and fluorescence, respectively, predictions of the fluorescent distribution can be made. The result clearly shows that cell length (C, dashed line) is a much better predictor of measured expression levels (C, blue area), than age (C, solid line). Also shown, is the distribution of expression levels obtained by stochastic simulation (C, gray line). Measured fluorescence distributions at (D) birth and (E) division (blue areas) are compared to stochastic simulations (gray lines). (F) Shows the comparison of the measured distribution of the fluorescence concentration of all extant cells (blue) and the simulations (gray line).

During balanced growth, concentrations of constitutively expressed proteins remain fixed; the cell’s content of a constitutively expressed protein therefore increases at the same rate as the cell volume. Our data clearly show that the total fluorescence levels (in arbitrary units) per cell correlate strongly with cell age (Fig. 4A and S5A) and cell length (Fig. 4B and S5B). This is to be expected, as cells that are older tend to be larger and are closer to having doubled their molecular content; the latter being a requirement for single cells under balanced growth. Although cell length and age strongly correlate, cell length explains more of the expression variability than age in both *B. subtilis* and *E. coli,* 79% and 73% vs 50% and 53% respectively. This indicates that the fluorescence signal (molecular content) of a cell is proportional to its volume (length in our case), as is expected for constitutive gene expression at balanced growth.

The relation between the mean fluorescence conditional on cell length, i.e. (*f |l_E_*), and cell length of extant cells (Fig. 4A and S5A), *l_E_*, can be transformed into the fluorescence distribution of the extant cell population when we take into account the cell age distribution and the dependency of *l_E_* on cell age. This gives rise to a good correspondence with the measured fluorescence distribution (Fig. 4C and S5C), indicating that the gene-expression noise – independent of cell age and cell length ‑averages out, due to a constant expression noise during the cell cycle (Fig. S10B and D). The gene-expression noise is best captured by the noise in the fluorescence concentration (fluorescence per fixed cell area), which is shown in Figs. 4F and S5F. The coefficient of variation in the fluorescence concentration equals 13% for *B. subtilis* and 14% for *E. coli.* Taken together, the fluorescence data indicate that more than 50% of the noise in total cell fluorescence originates from volume variability.

To test whether our algorithm accurately predicts cell growth and fluorescence distributions of populations at balanced growth, we performed stochastic simulations (see the Supplemental Information for details). As input we used the division length distribution (Ψ*_D_(l),* Fig. 2C), the daughter-over-mother volume ratio (*ρ*(*r*), Fig. 2B), and the measured growth rate (*k*). The expression model only accounts for constitutive zero-order synthesis of protein and protein dilution into new cells (no degradation). The parameters of the stochastic model of protein expression were obtained by linear fitting of the mean fluorescence at a given length (from Fig. 4B). To relate the measured protein fluorescence value to the protein copy number per cell used in the simulation, we use an arbitrary conversion constant. All stochastic simulation parameters can be found in the StochPy scripts available as supplemental material (S8).

The correspondence between simulated and measured distributions, shown in Figs. 2, 4, S4 and S5, indicates that the algorithm accurately recovers the statistical relations inherent in our experimental population.

It is interesting to note that the generation time distribution is used to calculate the average growth rate, however, its variability is not used as an input in the simulation algorithm. This implies that the variability observed in the generation time can be fully explained by the variability in the added volume per generation, ΔL. The overestimation in the fraction of large cells in the extant population (Fig. 2E) is propagated to the fluorescence distribution (Fig. 4C). Additionally, we observe a slight offset in the fluorescence distributions at cell birth and division (Fig. 4D-E). This shift is expected since our experimental setuρ recorded fluorescence with a resolution of ten minutes (whereas length is recorded every minute), leading to a shift between simulated and measured distributions. Fluorescence values are acquired, on average, 5 minutes after birth or 5 minutes prior to division, respectively. However, when the fluorescence concentration (Fig. 4E) for the experiment and the simulation is compared, we find a near perfect match.

## Discussion

We have validated a microscopic growth theory of bacterial cell growth with experimental data, illustrated how this theory can be exploited in the inference of probability distributions of single-cell growth characteristics, and employed the theory in the development of a fast and accurate stochastic simulation algorithm of the dynamics of a molecular circuit inside a growing single cell. This paper therefore contributes a quantitative methodology to single-cell physiology, a field that is in rapid development.

The microscopic theory relates the probability distributions of growth characteristics of single cells, such as cell sizes at birth and division, the generation time of cells, the growth rate of the cell population, the distribution of cell ages and sizes in the entire population of cells. Developed during the 1950-70s^27–29^, the theory describes the stochasticity of the growth of symmetrically dividing single cells, under the assumption that the cell population is at balanced growth. By comparison with real-time imaging of *E. coli* and *B. subtilis* growth, we show that the microscopic theory of cell growth has stood the test of time, and provides robust descriptions of the statistics of single-cell growth.

The microscopic growth theory has several applications. For example, it provides a robust way to check the consistency of experimental data assumed to be obtained at balanced cell growth. This would involve carrying out the same procedure as outlined in Fig. 2. Essentially, predicted distributions can be compared with measured ones. In this way, one not only checks whether cells were growing balanced, but also whether the algorithms used for time-lapse analyses are accurate. The theory could therefore also be conceived of as a component of a benchmarking protocol of software for microscopy-based time-lapse analysis of single-cell growth and fluorescent protein expression. Another application of the theory is to exploit it to estimate distributions of growth characteristics that were not measured, as we outlined in the results section.

While the microscopic growth theory describes the stochasticity of the growth of symmetrically dividing single cells, it does not deal explicitly with the stochasticity of molecular processes inside cells. Although pioneering studies on molecular networks followed shortly after^40^, the coupling between those two types of theories is a much more recent development (e.g.^6,22,41,42)^. Current analytical theories of the two-way coupling between single-cell growth and molecular circuit dynamics (e.g.^22,40,42)^, generally deal with small circuits. Numerical simulations are therefore indispensable for making predictions in single-cell physiology. Here we provide a novel demonstration of the microscopic theory as a means to speed uρ stochastic simulations of the growth of a single cell and of the dynamics of complex molecular circuits inside it.

Our algorithm simulates only a single lineage, from which we can retrieve the statistics of the entire lineage tree, which is what one generally obtains with real-time imaging of single cell growth. The advantage of this algorithm is that it is fast and in agreement with the microscopic theory of single cell growth. We validated the performance of this algorithm on experimental data and found that it is performing accurately. The current limitation of this simulation algorithm is that the molecular circuit cannot influence the growth behaviour of single cells. It simulates the dynamics of a molecular circuit on which we impose the stochasticity of single cells, which grow balanced, according to the microscopic growth theory, implemented in StOChPy^23^.

Even though the coupled stochastic simulation of growing cells and molecular circuits has been done (e.g.^41^ and^5^), we are not aware of any software package for stochastic simulation that has this capability. StOChPy is a flexible package, coded in the Python programming language. StOChPy has basic stochastic simulation algorithms (i.e. of the ‘Gillespie-type’), is readily extendible by the user, uses command-line instructions, allows for coding and saving of models in scripts, has a suite of statistical analysis and plotting tools, is compliant with SBML and can exchange models with the multi-purpose, deterministic modeling software package PySCeS^43^ (http://pysces.sourceforge.net) for systems biology.

Single-cell physiology has a big impact on cell biology. Its main study focus is the heterogeneity of isogenic cell populations, due to stochasticity at the level of growing cells and their molecular circuits, its origins and physiological consequences. The stochasticity of the growth and molecular circuits of single cells, and the nonlinear effects in those circuits, makes predicting the behaviour of single cells very complicated. We therefore have to rely on theory and simulations to interpret and predict the outcome of experiments that measure the physiology of single cells. This paper contributes some of the methodology required for a quantitative and predictive single-cell physiology.

## Methods

### Microscopy experiments

#### Strain, medium and culturing

*Escherichia coli* MG1655 derived MUK21 (see^44^ for details) (kindly provided by D. Kiviet), containing a genome integrated GFP gene under the control of the wild-type lac promoter, was revived from glycerol stock by inoculating directly into M9 minimal medium (42.2 mM Na_2_HPO_4_, 22 mM KH_2_PO_4_, 8.5 mM NaCl, 11.3 mM (NH_4_)_2_SO_4_, 2.0 mM MgSO4, 0.1 mM CaCl_2_), supplemented with trace elements (63 *μ*M ZnSO_4_, 70 *μ*M CuCl_2_, 71 *μ*M MnSO_4_, 76 *μ*M CoCl_2_, 0.6 *μ*M FeCl_3_), 0.2 mM uracil (all chemicals from Sigma) and 1 mM glucose as carbon source (M9-Glu). At intervals of 3 hours, the pre-culture was transferred twice to fresh M9 containing 10 mM lactose as carbon source (M9-Lac) and 1 mM of IPTG (M9-Lac-I), before inoculating an overnight culture to a final optical density (OD, 600 nm) of 2.5 × 10^-7^ in M9-Lac-I. After 16 hours, the culture was again diluted to an OD600 of 0.0025. When the culture reached an OD600 of 0.01, 2μL was transferred to a 1.5% low melt agarose pad freshly prepared with M9-Lac-I.

*Bacillus subtilis* strain B1115 (amyE::Phyper-spank-sfGFP, Spcr) was constructed from parental strain BSB1 168 *trp+* using pDR111 (kindly provided by D. Rudner). B1115 was revived from glycerol stocks by directly inoculating into MM minimal medium (40 mM MOPS, 1.23 mM K_2_HPO_4_, 0.77 mM KH_2_PO_4_, 15 mM (NH_4_)_2_SO_4_, 0.8 mM MgSO_4_), supplemented with trace elements (80 nM MnCl_2_, 5 *μ*M FeCl_3_, 10 nM ZnCl_2_, 30 nM CoCl_2_, 10 nM CuSO_4_), 5 mM glucose and 1 mM IPTG (MM-Glu-I) to a final OD of 4.5 × 10^-7^. After incubation for 16 hours, the culture was diluted in TSS minimal medium (37.4 mM NH_4_Cl, 1.5 mM K_2_HPO_4_, 49.5 mM TRIS, 1mM MgSO_4_, 0.004% FeCl_3_, 0.004% Na_3_-citrate.2H_2_O), supplemented with trace elements, 5 mM glucose and 1 mM IPTG (TSS-Glu-I) to an OD of 0.001. When the culture reached an OD600 of 0.02, 2 UL was transferred to a 1.5% agarose pad freshly prepared with TSS-Glu-I.

All cultures were incubated at 37° C, shaking at 200 rpm. Once seeded with cells, agarose pads were inverted and placed onto a glass bottom microwell dish (35 mm dish, 14 mm microwell, No. 1.5 coverglass) (Matek, USA), which was sealed with parafilm and immediately taken to the microscope for time-lapse imaging.

#### Microscopy and data analysis

Imaging was performed with a Nikon Ti-E inverted microscope (Nikon, Japan) equipped with 100X oil objective (Nikon, CFI Plan Apo *λ* NA 1.45 WD 0.13), Zyla 5.5 sCmos camera (Andor, UK), brightfield LED light source (CoolLED pE‑ 100),fluorescence LED light source (Lumencor, SOLA light engine), GFP filter set (Nikon Epi-Fl Filter Cube GFP-B), computer controlled shutters, automated stage and incubation chamber for temperature control. Temperature was set to 37° C at least three hours prior to starting an experiment. Nikon NIS-Elements AR software was used to control the microscope.

Brightfield images (80 ms exposure time at 3.2% power) were acquired every minute for 8 hours. GFP fluorescence images (1 second exposure at 25% power) were acquired every 10 min. Time-lapse data were processed with custom MATLAB functions developed within our group. Briefly, an automated pipeline segmented every image, identifying individual cells and calculating their spatial features. Cells were assigned unique identifiers and were tracked in time, allowing for the calculation of time-dependent properties including cell ages, cel sizes (areas and lengths), elongation rates and generation times. In addition, the genealogy of every cell was recorded.

Only cells from within the exponential population growth phase (see Fig. S1) were considered in the analysis. For cells expressing a fluorescent construct, the fluorescence at birth and division were taken as the first and last measurement during the single cell time trace. Because we measured the GFP fluorescence once every ten minutes, an average deviation of five minutes from *a =* 0 and *a = T* is present in the fluorescence data.

### Stochastic simulations

Stochastic simulations were done with the direct method algorithm^45^ extended with cell growth and division, as implemented in the CellDivision module of the stochastic simulation software package StochPy^23^. Each stochastic simulation was continued for 10^4^ generations. More information on installing and using StochPy can be found in the *StochPy User’s Guide* which together with additional example sessions is available online at http://stochpy.sf.net.

## Acknowledgements

FJB acknowledges funding from the Netherlands Research Organisation (NWO-VIDI 864.11.011). We thank Maxim Moinat for extending the StochPy simulation package and Coco van Boxtel for initial fluorescence microscopy experiments and testing the image analysis, cell segmentation and tracking algorithms.

## Author contributions statement

JvH and FJB conceived the experiments. JvH and NN conducted the experiments. JvH, MK, AD, TM and FJB developed software and algorithms. JvH, MK and FJB analysed data. JvH, MK and FJB wrote the manuscript. All authors reviewed the manuscript.

## Competing financial interests statement

We do not have any competing financial interests to declare.

## Supporting Information

Contents

References 11

1 Confirmation of balanced growth during the microscopy experiment

2 Cell length and fluorescence data of a balanced growing ***E. coli*** population

3 The cell-size homeostasis mechanism as evident from our data

4 Cell length as an approximation for cell volume

5 Growth rate of ***B. subtilis*** and ***E. coli*** along the cell cycle, in terms of cell length and cell age

6 Fluorescence concentration and fluorescence noise for ***B. subtilis*** and ***E. coli*** along the cell cycle, in terms of cell age

7 StochPY extended with cell growth and division

7.1 Stochastic simulation of single-cell growth and gene expression matches theory

7.2 Conditions to interrelate a lineage and full tree simulation

7.3 The joint distribution of cell age and generation time

7.4 Illustrations of StochPy simulations with cell growth and division

7.5 Comparing StochPy simulations with cell growth and division to analytical solutions

Case Study 1: Volume statistics are independent of the model • Analytical solutions: Case Study 1 Volume distributions • Case Study 2: mRNA synthesis • Analytical solutions: Case Study 2 mRNA synthesis • Poisson distributed molecule copy numbers at a specific age • mRNA copy number distribution of a sample of extant cells • Case Study 3: mRNA synthesis and degradation • Analytical solutions: Case Study 3 mRNA synthesis and degradation • Poisson distributed molecule copy numbers at a specific age • Molecule copy-number distribution of a sample of extant cells

### 1 Confirmation of balanced growth during the microscopy experiment

During steady state balanced growth, the specific growth rate and other properties of the population should remain fixed for a duration longer than several generation times^27^. We confirmed this by plotting the logarithm of the total cell length of the population as function of time and identifying a linear region, during which balanced growth occurs (Fig. S1).

**Figure S1.**
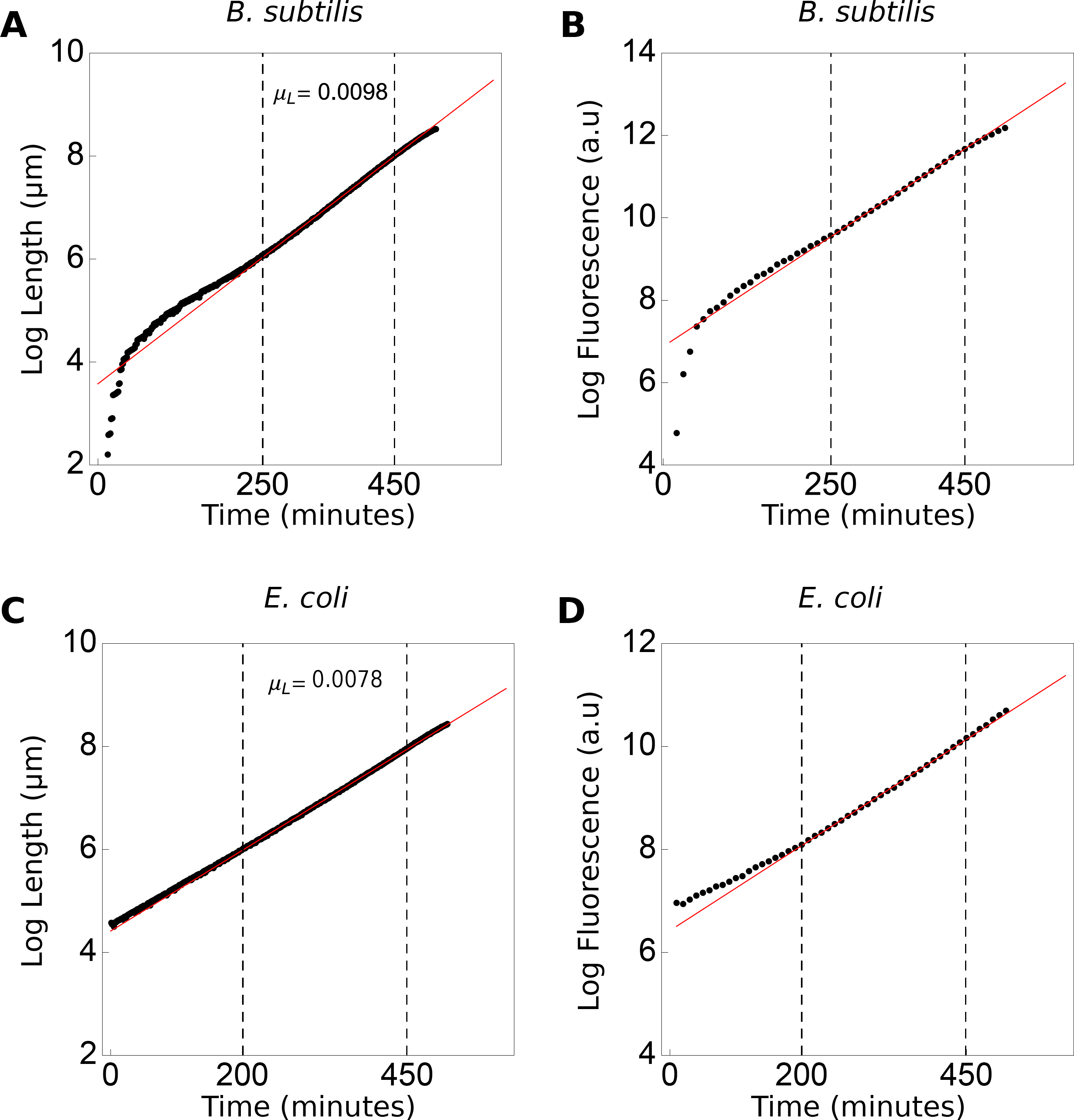
Linear fit of the log transformed total cell length and fluorescence of the *B. subtilis* and *E. coli* population. The total length of all individuals in the (A) *B. subtilis* population increases with 0.0098 *min*^-1^ and for *E. coli* this rate is 0.0078 *min*^-1^ (C). The dashed lines marks the region of balanced growth, i.e. where growth rate is fixed over multiple generations, used for data analyses. Similarly, the rate of total fluorescence increase is fixed during balanced growth, for both (B) *B. subtilis* (D) and *E. coli*.

During balanced growth, the total fluorescence of cells, which corresponds to the levels of a constitutively expressed protein, should increase exponentially with time (Fig. S1). This indicates that the protein abundance changes in proportion to the total cell volume, which in our case simplifies to proportional increase with cell length, as the cell maintain a fixed cell width (Fig. S2). That the concentration of a constitutively expressed protein remains fixed during balanced growth can be deduced from the differential equation of the concentration, denoted by *c*, which equals the ratio of the protein copy number *n* (assumed proportional to protein fluorescence) and the cell volume, *V*,

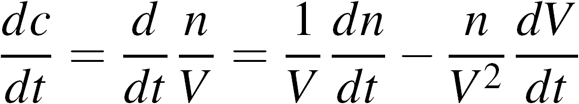

When concentration is at steady state then:

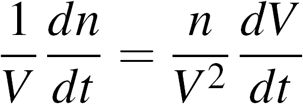

We can now identify the specific growth rate from,

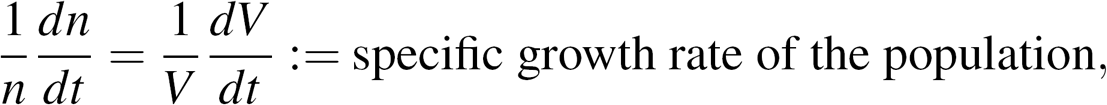

indicating that *n* and *V* increase exponentially in time. From this we can conclude that during exponential growth the concentration of the protein remains fixed, this implies that the protein copy number increases exponentially at the same rate as the cell volume.

Another requirement of balanced growth is that the probability distributions of growth characteristics are invariant with time, which is illustrated by Fig. S3.

**Figure S2.**
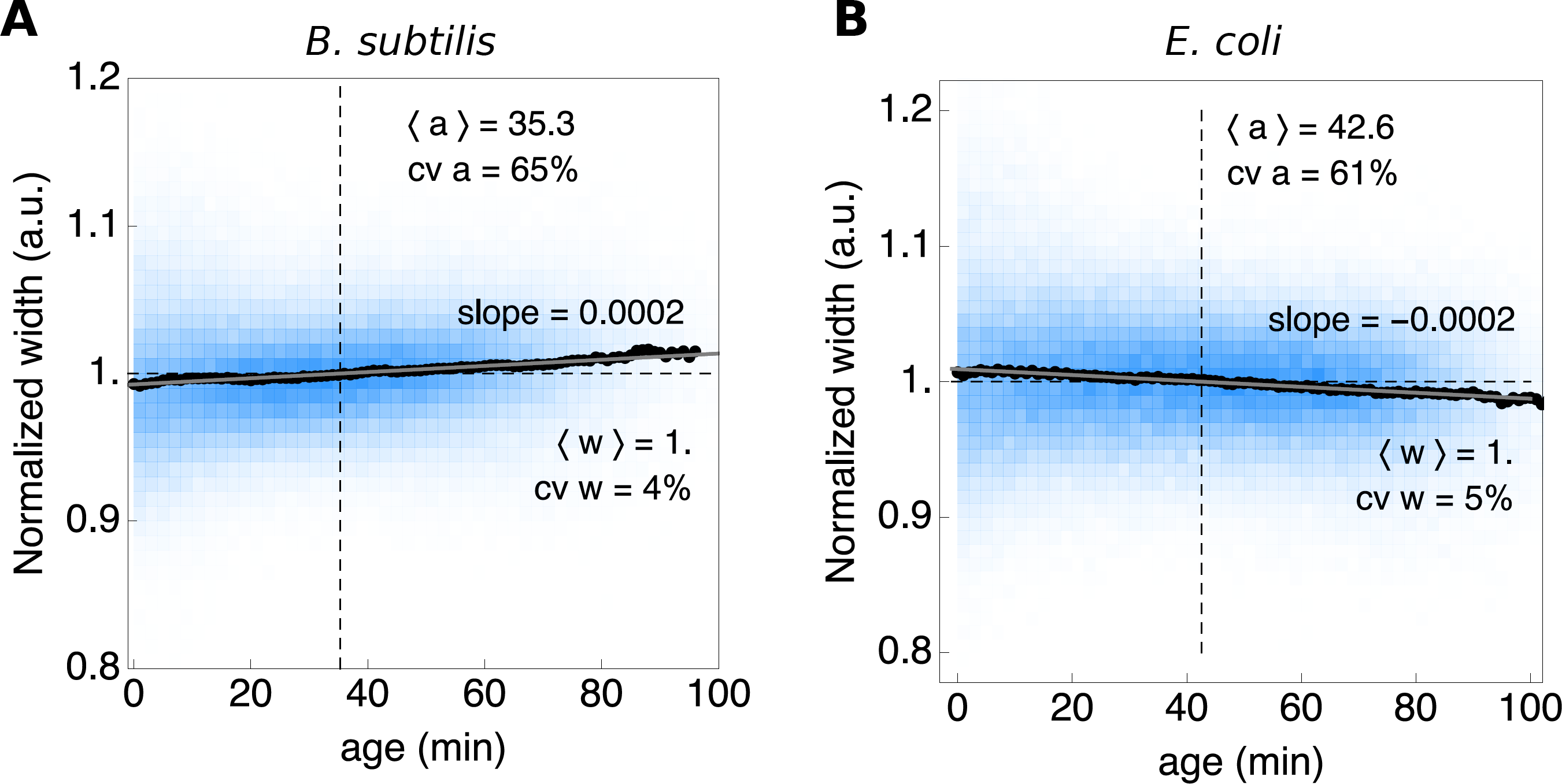
Cell width as function of age for a B. subtilis and E. coli population. The width of cells remain approximately constant during their life cycle.

### 2 Cell length and fluorescence data of a balanced growing *E. coli* population

In addition to the *B. subtilis* data, which we presented in the main text, we measured cell growth characteristics (Fig. S4) and gene expression (Fig. S5) in a balanced-growing *E. coli* population. As for *B. subtilis,* the *E. coli* experimental data shows a remarkable agreement with the microscopic growth theory and stochastic simulations.

### 3 The cell-size homeostasis mechanism as evident from our data

A single-cell growth characteristic that has since recently received a lot of attention is the mechanisms which cells use to achieve size homeostasis during balanced growth^5^. Since the probability distributions for cell size are constant during balanced growth, cells that are either smaller or larger than the average cells size compensate for their size discrepancies. At least three mechanisms have been proposed that lead to cell-size homeostasis^5^. Cells can either be ‘sizers’, ‘adders’ or ‘timers’, or they follow one of two mixed mechanisms: ‘sizer-like adder’ or ‘timer-like adder’^39^.

Sizer, adder and timer mechanisms can be distinguished from the slope of the relation between the average length that cell adds during a single generation and its birth length (i.e. (Δ*l*|*l_B_*) as function of *l_B_*)^5,39,46^. The slopes we find in the (Δ*l*|*l_B_*)-vs-*l_B_* plot (Fig. S6) are slightly negative (≈ −0.3), indicating that both *B. subtilis* and *E. coli* follow a sizer-like adder mechanism.

### 4 Cell length as an approximation for cell volume

During balanced growth, cell volume (and cell length when the width remains fixed) double each cell cycle. On average, the mean volume at division is twice the mean birth volume. We measure cell growth with a time resolution of one minute, the determined moment of cell birth and cell division deviates on average by half a minute. Within this time window minor cell growth and deformation changes might occur, disturbing the expected 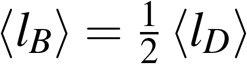 relation. Indeed, the observed cell length which we use as a measure for cell volume, does not double exactly in length during a cell cycle. When we use the combined length of both daughter cells as measure for division length the expected 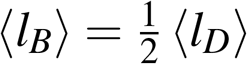 relation holds.

Since the deviation in cell length is more than expected in a minute of cells growth, we consider the influence of rod-shaped bacterial growth to bring forward a possible explanation. When the exact moment of cell division is not determined perfectly, the mother cell might represent more of a single rod shape rather than two rods (Fig. S8). This figure indicates that a single cell can double its volume while not doubling its length.

To estimate the maximal deviation between volume and length growth, we consider cells as perfect rod-shaped until the moment of division. We observed a constant cell width during a cell cycle (Fig. S2), hence the radius of the cell is fixed. The difference in cell length and volume growth is caused by the cell poles, which are rounded. The in length increase to double the volume of a rod is *2L_b_ – α*, in which *α* is given by:

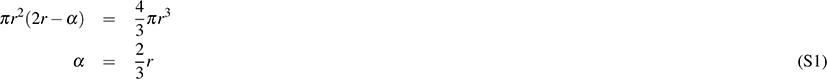

this solves the cylindrical height to equalise the volume of a sphere (the sum of the two rounded poles) to a cylinder. This means that the deviation in observed cell length growth can be at max 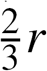. In table S1 we provide this minimal estimate of cell length compared to the measured cell length for *L_B_, L_D_,* and Δ*L*. The measured mean lengths fall somewhere between exact doubling of cell length and the calculated length increase based on a rod-shaped cell throughout the entire cell cycle. Given the length statistics of *B. subtilis* we predicts a minimal Δ*L* of 2.19 *μ*m. For the *E. coli* data the predicted minimal ΔL is 1.43 *μ*m. It is known that *B. subtilis* and *E. coli* divide using different division modes^47^. In *E. coli* the cell membrane grows inwards while *B. subtilis* assembles a septal wall. Due to this differences in division mechanisms, the ΔL of *E. coli* might be more close to *L_b_.* While the Δ*L* of *B. subtilis* is more close to discussed rod-shaped mother cells. Of interest, using the combined length of both daughter cells as measure for division length, gives (Δ*L*) = 2.50 for *B. subtilis* and (Δ*L*) = 1.62 for *E. coli.*

**Figure S3.**
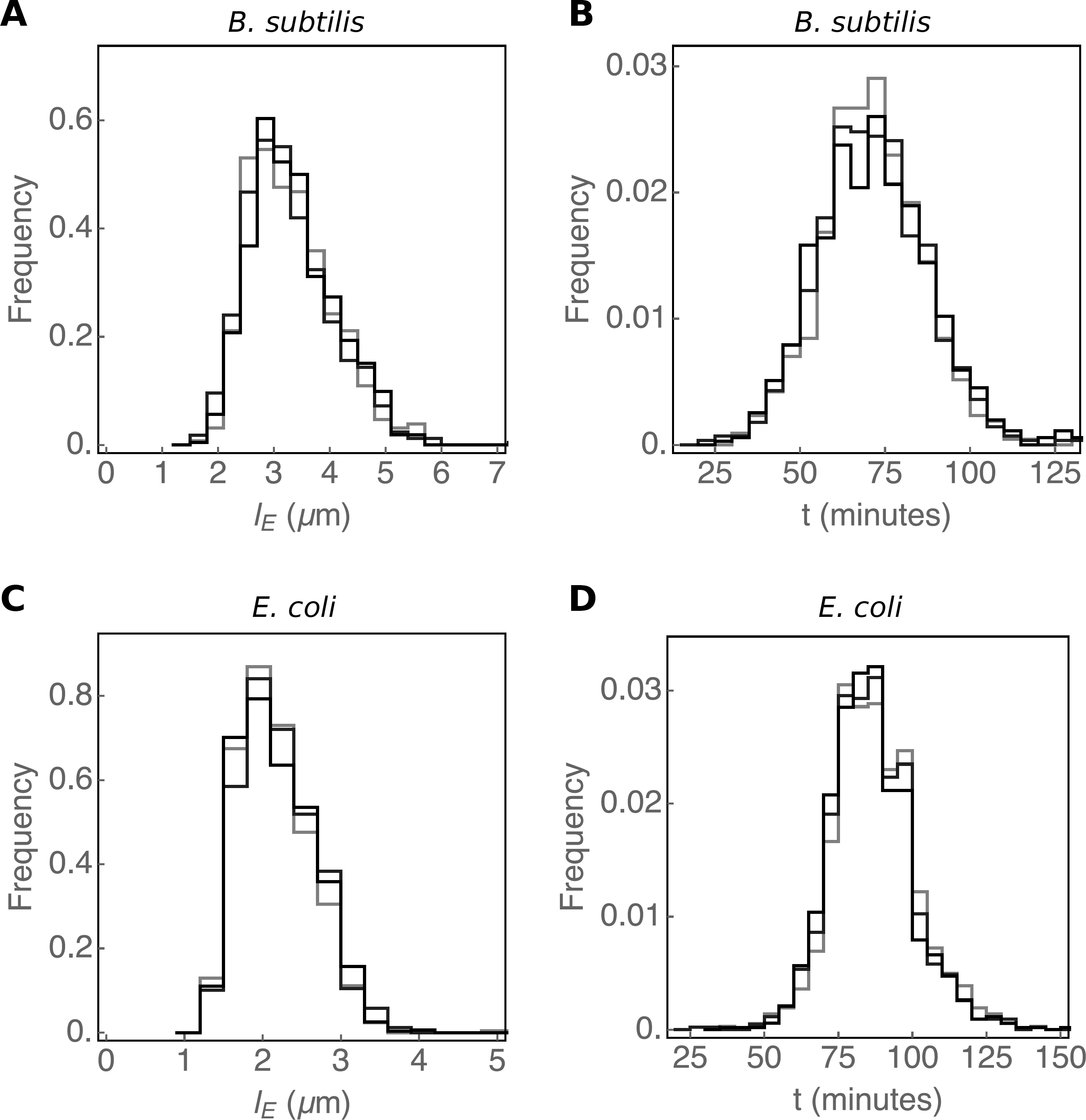
Balanced growth is characterised by the time-invariant distribution of growth characteristics. A defining characteristic of balanced growth is that the probabilities to observe cells with particular growth properties – their phenotype – are fixed and the associated probability distributions are therefore time invariant. The extant cell length (A and C) and generation time (B and D) distributions sampled at three different time points during balanced exponential growth (see Fig. S1) are shown for a *B. subtilis* and *E. coli* population.

**Figure S4.**
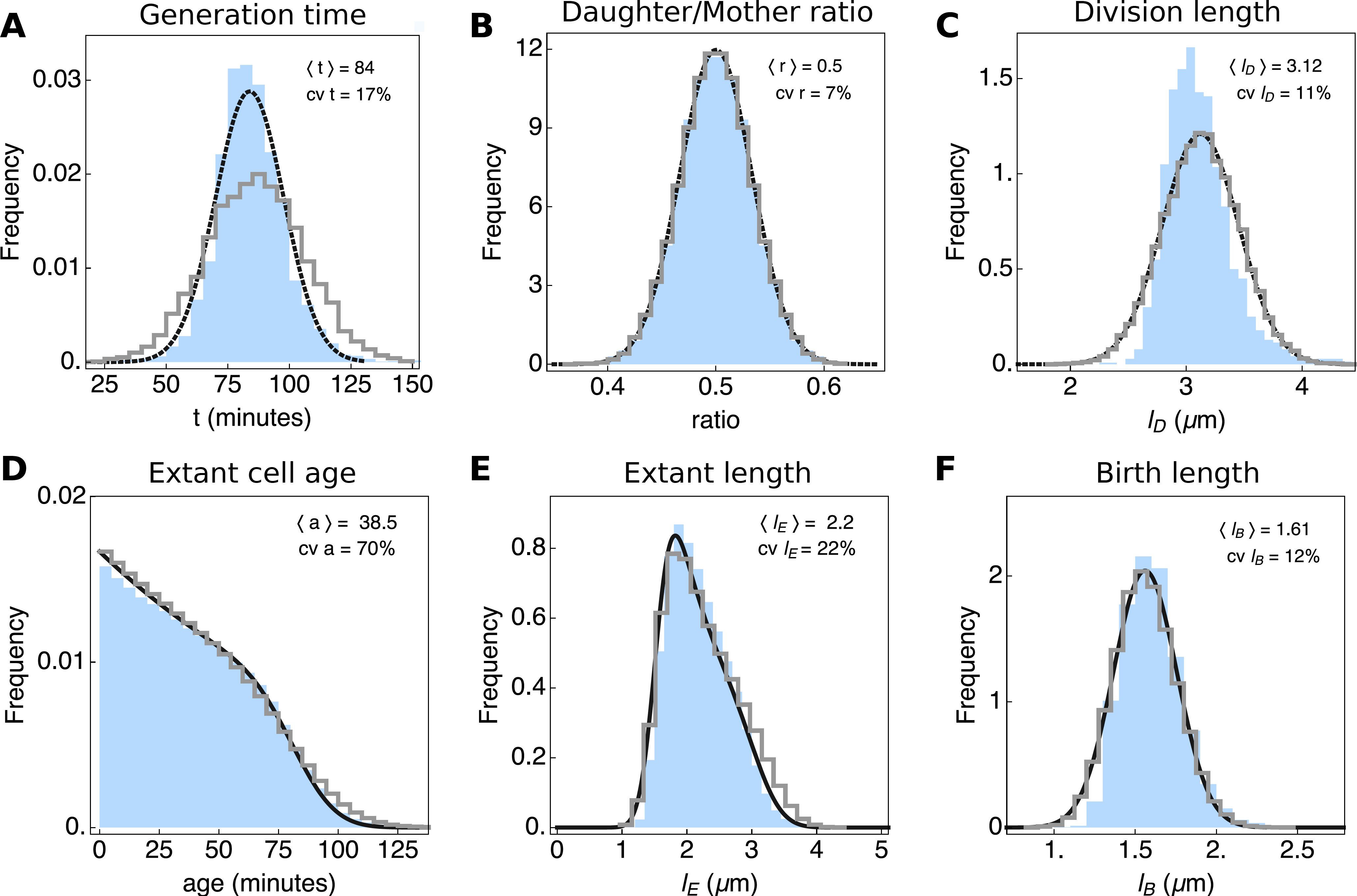
Growth characteristics of *E. coli*. Shown are results comparing the microscopic growth theory relations derived by Collins & Richmond^29^, Powell^28^ and Painter & Marr^27^ and experimental single-cell growth data. This figure is the same as Fig. 2, but shows data for *E. coli.* The probability distributions obtained from experimental data are shown in blue, the predicted distributions (obtained using the relations defined in eqs. 1 – 4 in Fig. 2), are shown in black in (D), (E) and (F), and the results of our simulation algorithm are shown as grey histograms. In (A), (B) and (C), black curves indicate fits to the measured data. The sample sizes for the experimental data are 4558 extant cells, 3617 cells at birth and 1867 cells at division.

**Table S1.**
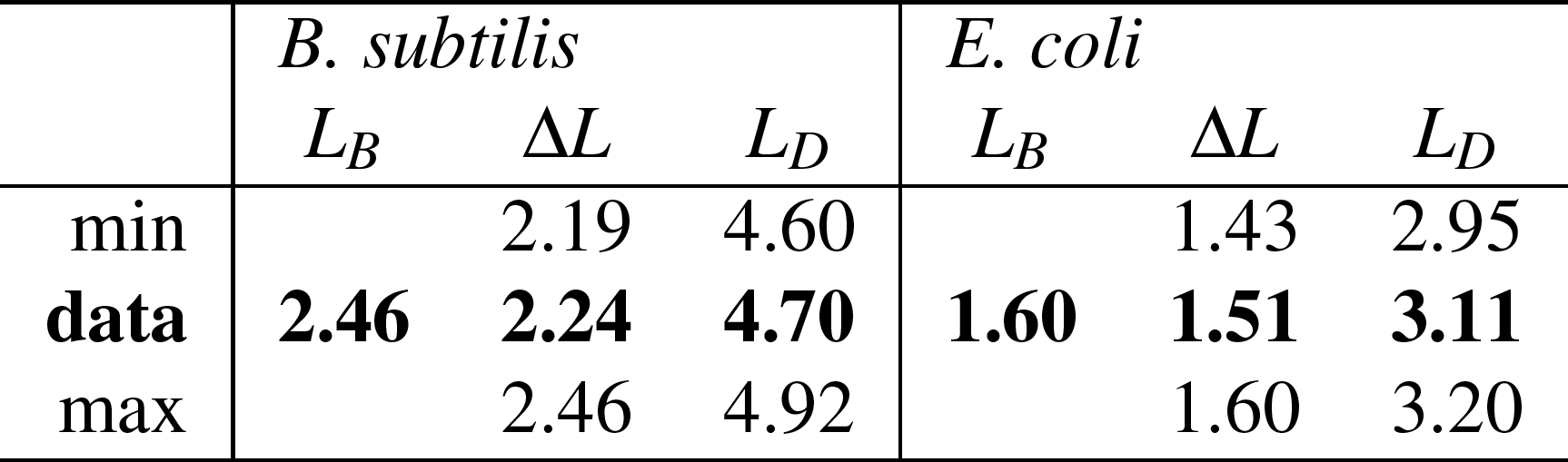
Average cell length calculated from the birth length distribution (ℓ_B_). The range is given by the perfect rod-shape model and the double of birth length. The data indicates that the measured length falls within the expected range.

|  | *B. subtilis* |  |  | *E. coli* |  |  |
| --- | --- | --- | --- | --- | --- | --- |
| | $L_B$ | $\Delta L$ | $L_D$ | $L_B$ | $\Delta L$ | $L_D$ |
| min |  | 2.19 | 4.60 |  | 1.43 | 2.95 |
| **data** | **2.46** | **2.24** | **4.70** | **1.60** | **1.51** | **3.11** |
| max |  | 2.46 | 4.92 |  | 1.60 | 3.20 |

### 5 Growth rate of *B. subtilis* and *E. coli* along the cell cycle, in terms of cell length and cell age

Figure S9 shows the instantaneous growth rate of *E. coli* and *B. subtilis* at particular progression extends along their cell cycle, at a particular length (Fig. S9A and C) and a particular age (Fig. S9B and D). The conditional mean growth rate, i.e. (μ|*l*) and (μ|*age*) is fixed along the cell cycle.

**Figure S5.**
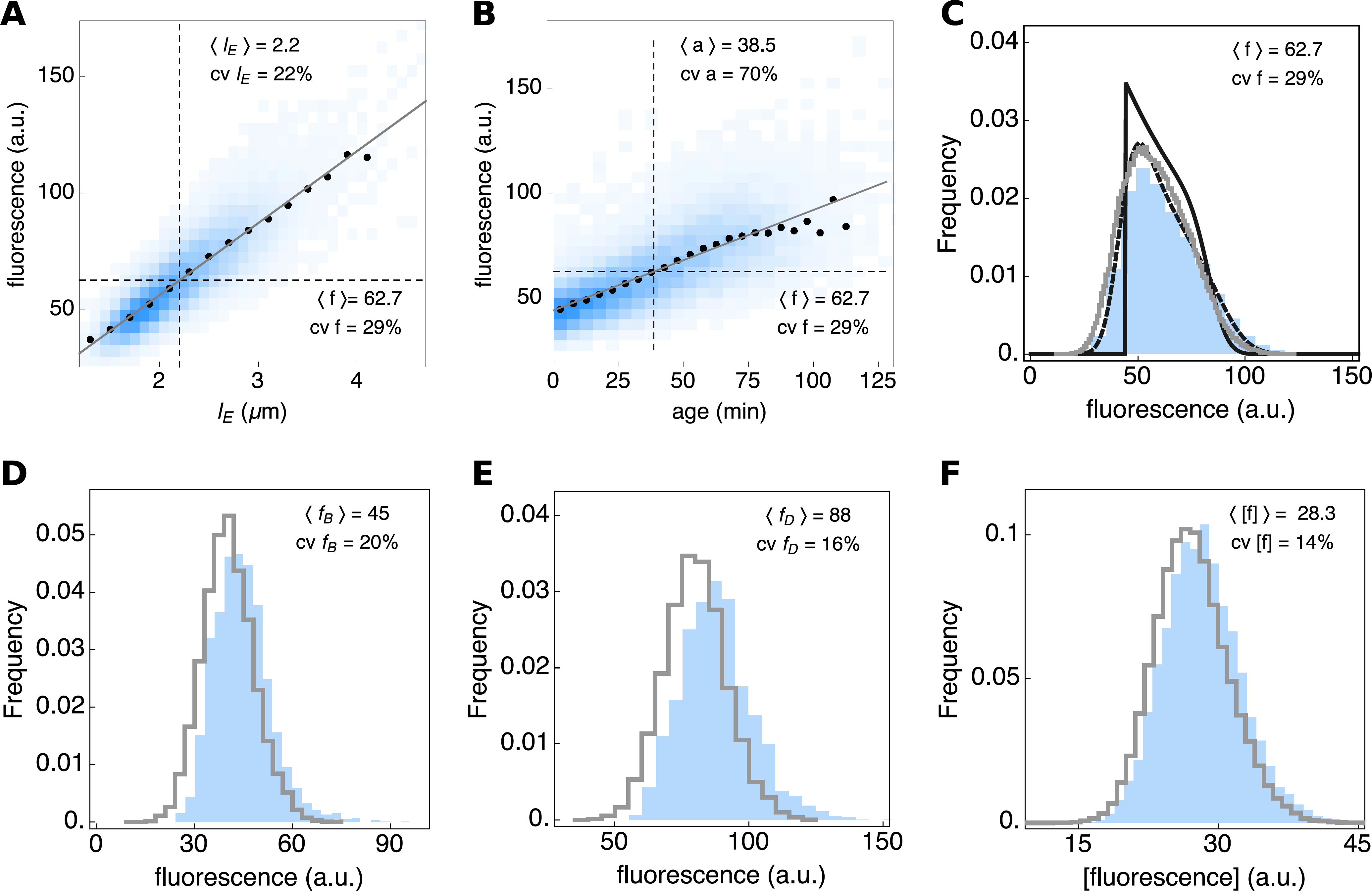
Expression data of the reporter gene for *E. coli.* (A) Whole-cell fluorescence as function of cell length with black dots as conditional mean <*f* |*l*_E_>. (B) Cell fluorescence as function of cell age with black dots as conditional mean *<f* |*age*>. (C) Whole-cell fluorescence probability density: experimental data (blue), predicted fluorescence probability density obtained from the extant length distribution and the relation between fluorescence and length (shown in A) (dashed, black line), predicted fluorescence probability density obtained from the age distribution and the relation between fluorescence and age (shown in B) (black line) and the stochastic simulation (grey line). The experimentally determined distributions (blue) of whole-cell fluorescence at (D) cell birth, (E) cell division and (F) the distribution of fluorescence concentration of extant cells, is compared to simulations (grey lines).

### 6 Fluorescence concentration and fluorescence noise for B. *subtilis* and E. *coli* along the cell cycle, in terms of cell age

The fluorescence concentration and noise hardly change as function of the progression along the cell cycle as is indicated by the slopes of the conditional mean lines as function of cell age in Fig. S10. Only the fluorescence noise data of *E. coli* shows a decrease of about 20% from birth to division.

**Figure S6.**
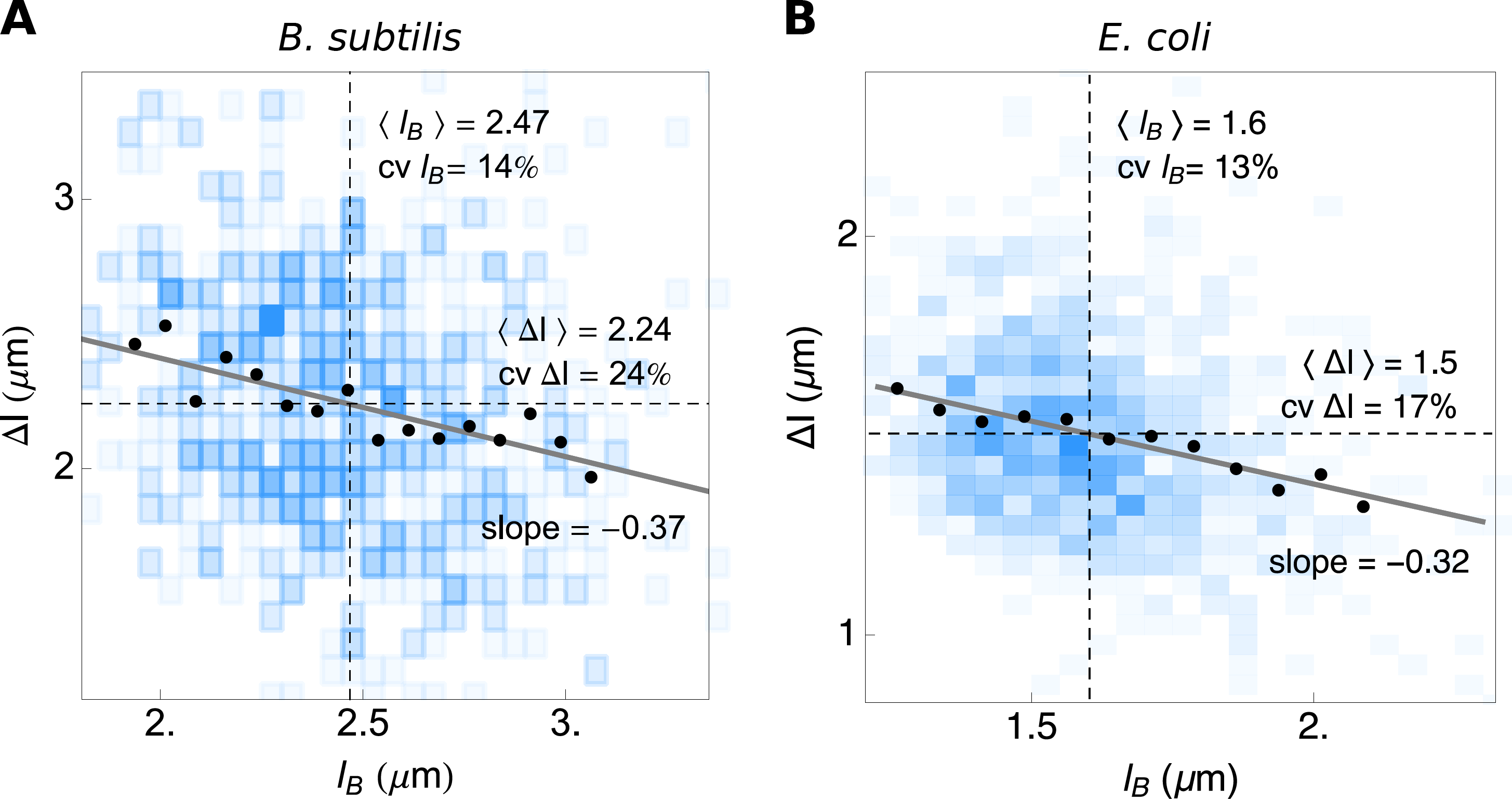
Cell-size homeostasis mechanisms inferred from the single-cell growth data of *B. subtilis* and *E. coli.* From the slope of the dependency of average cell length added during a single generation on the length at cell birth (<Δ*l*|*l_b_*>-vs-*l_b_*), the cell-size homeostasis mechanism can be inferred. The slopes of about −0.3 indicate that, under the conditions tested, (A) *B. subtilis* and (B) *E. coli* behave mostly as adders with a small sizer effect.

**Figure S7.**
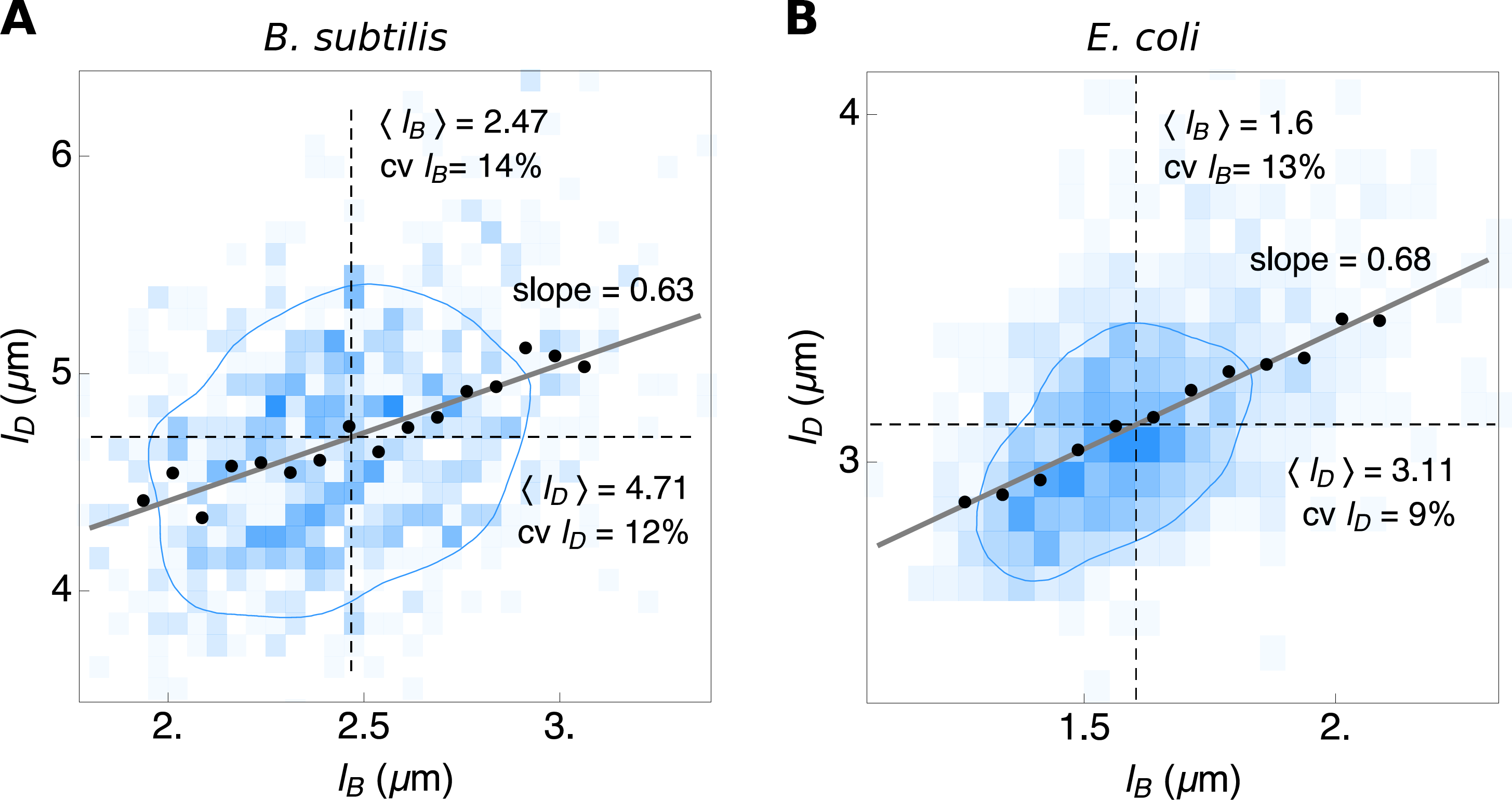
Relation between the cell volume at birth and division inferred from the single-cell growth data of *B. subtilis* and *E. coli.* The length at cell division as function of the length at birth (<*l*_D_|*l*_B_>-vs-*l*_B_) of (A) *B. subtilis* and (B) *E. coli* shows a weak positive correlation.

**Figure S8.**
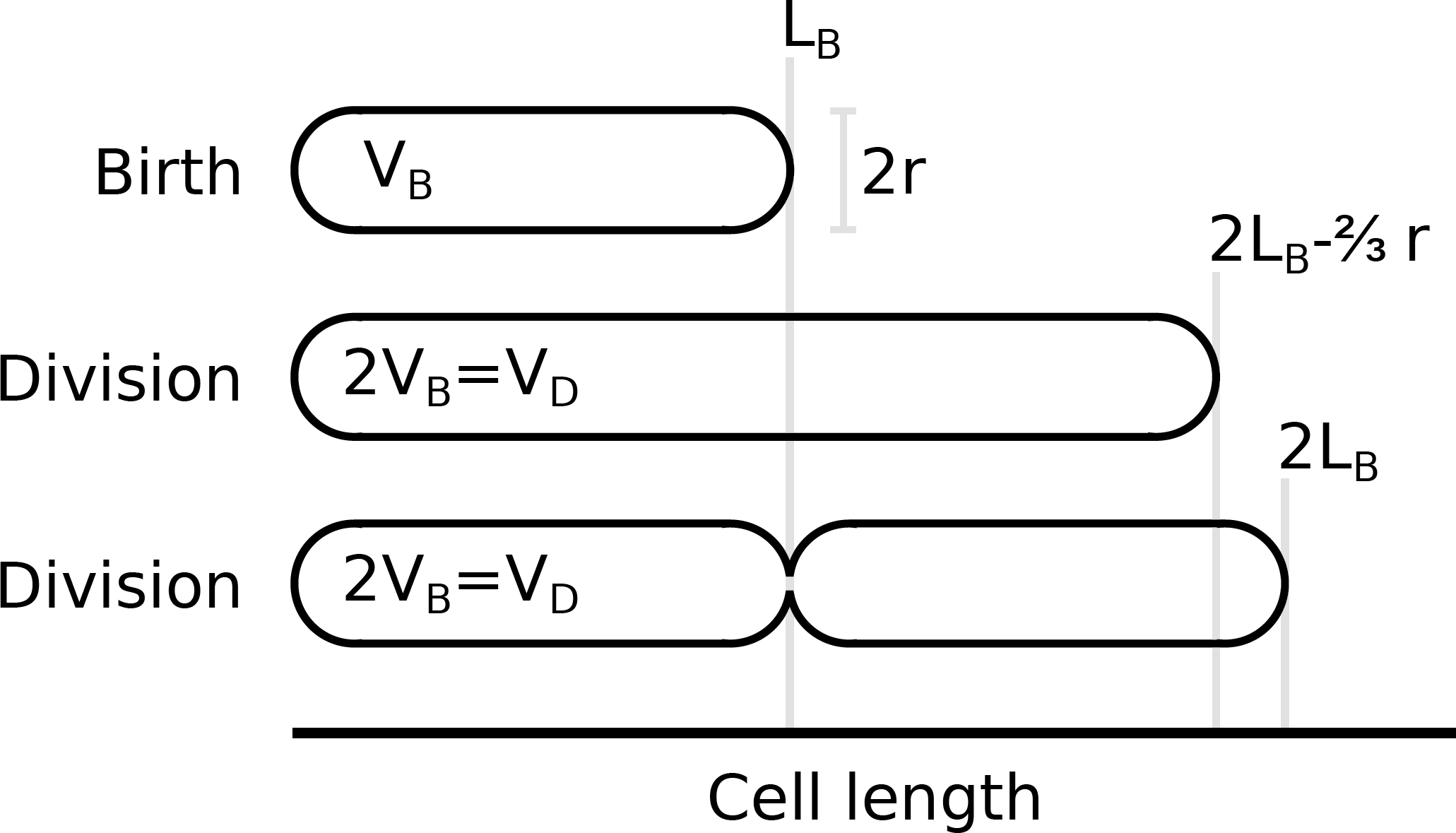
Cell shapes at birth and division. At cell division the average cell volume is doubled compared to the birth volume. Depending on the cell shape of the mother cell at division, the cells do not display full length doubling during a single cell cycle. The maximal deviation between volume and length increase can be calculated and is 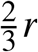.

**Figure S9.**
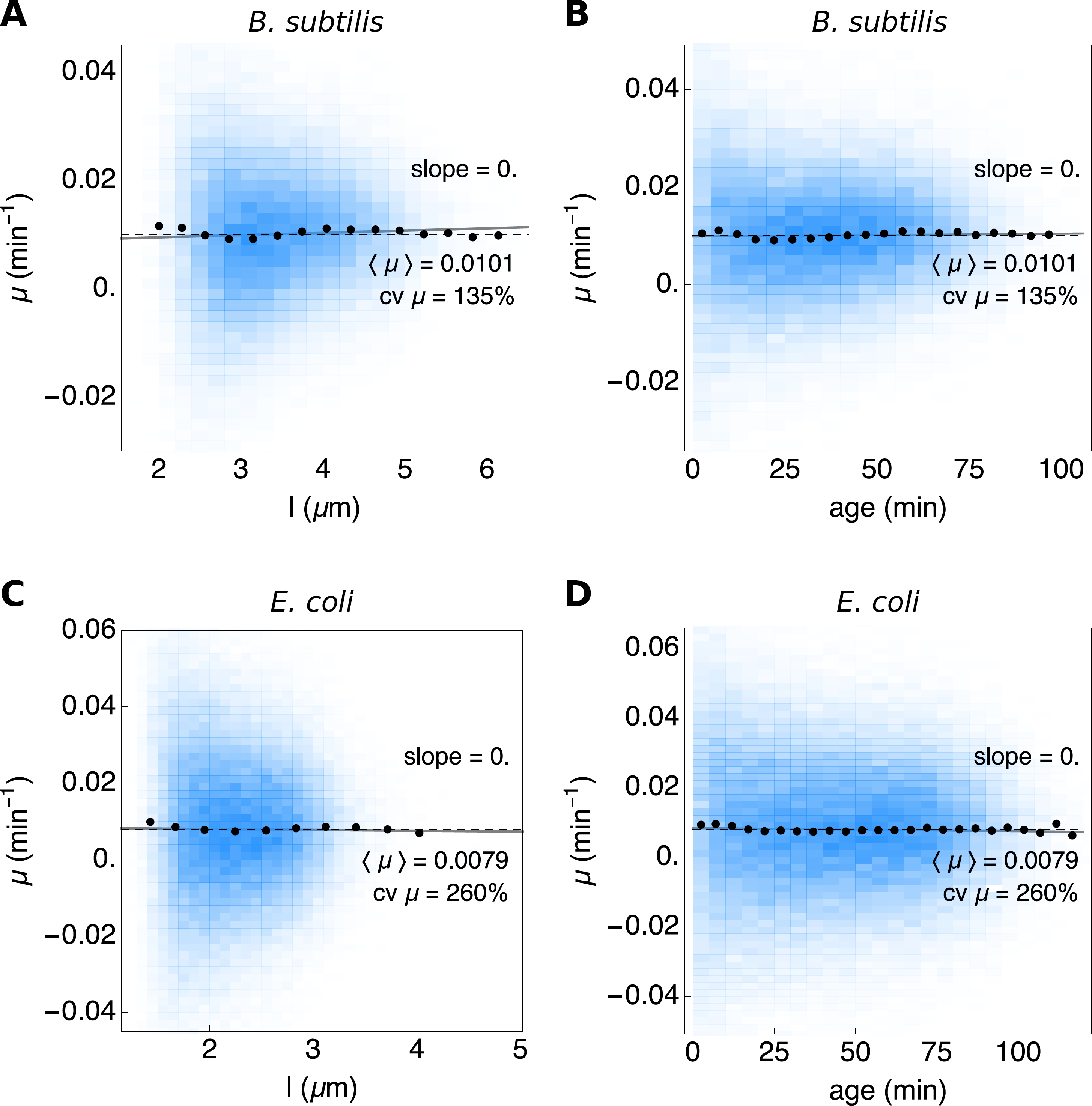
Instantaneous growth rate as function of age and cell length for *B. subtilis* and *E. coli.* The blue density histograms give the distribution of the instantaneous growth rate as function of length (A and C) and age (B and D). The population means are shown by the dashed black lines. A linear fit to the full dataset is shown in gray and the conditionals μ |*l* and μ |*age* by the black dots.

**Figure S10.**
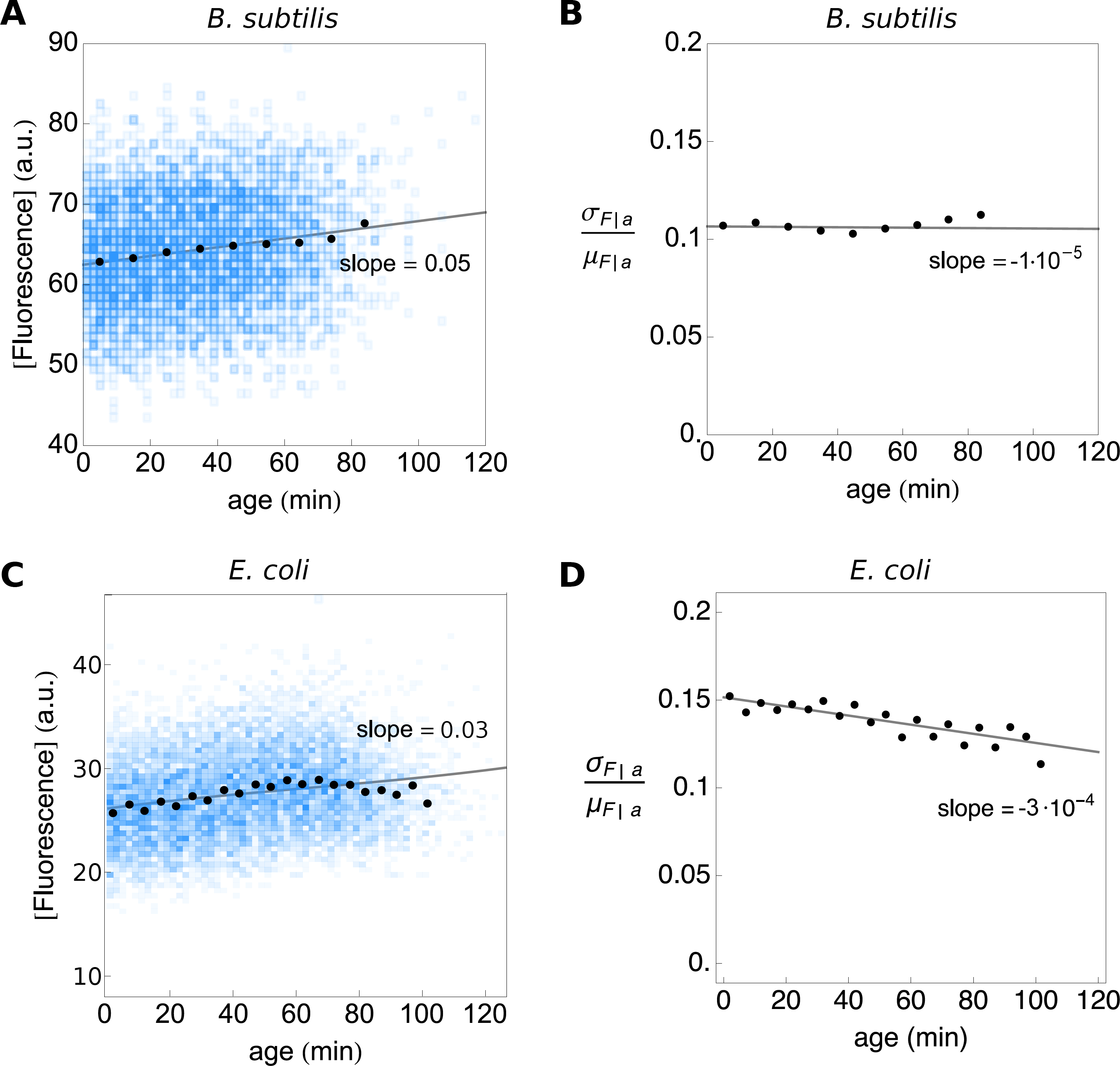
Concentration of fluorescence and fluorescence noise as function of age for the *B. subtilis* and *E. coli* data. The average concentration of fluorescence remains stable during a cell cycle. The cv of the concentration of fluorescence does not increase as function of age, explaining the similarity between the theoretically predicted and observed expression distribution (see Fig. 4C).

**Figure S11.**
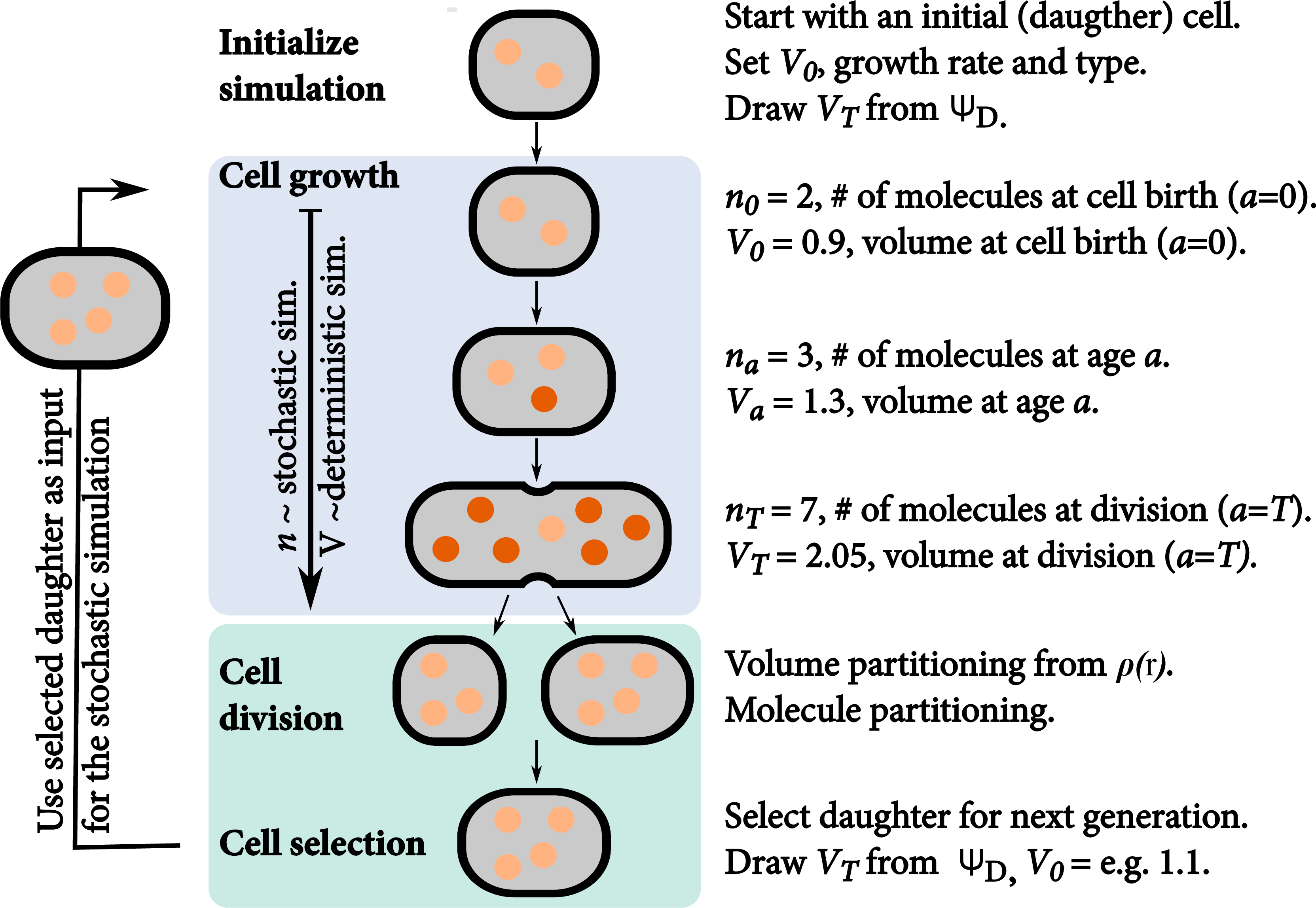
The SSA with cell growth and division as incorporated in StochPy. The simulation starts with one daughter cell of which we track a single lineage through time. We perform a stochastic simulation until the generation time *T* is reached. Then, the mother volume is partitioned between both daughter cells. Molecules are subsequently partitioned (volume dependent) between both daughters. The light molecules are inherited from the mother cell, darker ones are newly synthesised during the cell cycle. This procedure is repeated until either the number of generations is reached, the desired end time is reached, the desired number of time steps is reached, or all reactions are exhausted.

### 7 StochPY extended with cell growth and division

The stochastic simulation algorithm that couples the simulation of molecular circuits to that of cell growth and division is implemented in StochPy^23^ and available for download from http://stochpy.sf.net. Figure S11 gives an overview of how we extended the SSA with cell growth and cell division.

We discuss the steps outlined in Fig. S11 step-by-step (the notations used are explained in Table S2):

1. **Initialize simulation.** In the extended SSA, containing cell growth and division, we simulate a single lineage, so we start the simulation with a single cell. Before we can perform a simulation we have to initialize the simulation by setting various parameters. For instance, we have to set the cell volume of the initial cell (V_0_). A cell can be in any cell-cycle stage (0 ≤ *a ≤ T*) where *a* is the cell age and *T* the generation time—the time period between two consecutive cell divisions. We parameterise the initial cell to start at the beginning of the cell-cycle stage, i.e. *a =* 0. During the simulation, the cell grows deterministically, at the specified specific volume-growth rate (μ) and according to a particular type of growth—we support both exponential and linear volume growth rates. Next, we have to determine when the tracked cell divides. We do this by drawing a volume (V_T_) at which the mature mother cell divides into two daughter cells. The volume at which the mother cell divides is drawn from a probability density function (PDF) Ψ_D_. This PDF is independent from the volume at birth, which makes the volume at division independent from the volume at birth. Given values for V_0_, *V_T_*, u, and the growth, the generation time *T* can be calculated. We illustrate this here for an exponential growth rate:

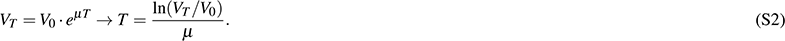
2. **SSA coupled to cell growth.** Once we know *T*, we can start the SSA until the division event at *a = T*. Modeling cell growth implies that *V* increases during the simulation from birth *(a =* 0) to division *(a = T*). In a growing cell, reacting molecules require more time to find each other, thus the reaction waiting times for these kind of reactions increases. This means that the propensity functions of diffusion limited reactions (i.e. second and higher-order reactions) depend on *V*. We therefore inserted *V* as a variable in the respective propensity function,

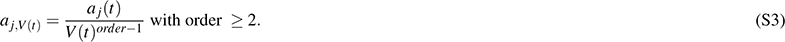

The propensity functions of zero and first-order reactions are unaffected by *V*. While we model *V* deterministically we only update *V* at each stochastic event in the simulation. This means that we calculate the time until the next reaction fires based on the *V* at the time of firing of the previous reaction; *V* is larger at the moment of reaction execution which results in a underestimation of the reaction time of second and higher-order reactions. This effect is negligible if the volume difference between the two consecutive firings, ΔV, is small. An alternative method could be to add volume as an additional reaction stochastic reaction that fires frequently as was done by^41^, but this slows down the simulation.
3. **Cell division.** The SSA continues until the generation time *T* is reached (and *V = V_T_*). Both *V_T_* and *x_T_* are then partitioned between the two daughter cells. The partitioning ratio is drawn from the PDF ρ(r). Daughter one and two receive a volume of V_*d1*_,_0_ = *r · V_T_* and V_*d2*_,_0_ = (1 – *r) · V_T_* respectively. The ρ*(r)* distribution should be symmetric around a mean of 0.5, otherwise a bias for one daughter is created. The partitioning of molecules between both daughter cells—which is done next—is also a stochastic process and depends on the cell volumes of both daughters. This partitioning of molecules is, therefore, modeled with a volume-dependent binomial distribution. More specifically, the probability that a specific molecule is inherited by daughter one is modeled as *V_d1,0_/V_T_*. This means that the number of molecules, with copy number *n*, inherited by daughter one can be drawn as a random sample from a binomial distribution with *n* number of trials and success probability V_*d1*,0_/V_T_. The process is repeated for each species. Not all cellular constituents should be binomially distributed between both daughter cells. DNA is an example of this; each daughter cells receives one copy of the chromosome in a normal cell division event. These kind of cellular constituents divide exact. Stochastic simulations also allow the definition of fixed species—species that do not change in copy number of concentration over time—which are not divided during a cell division event.
4. **Cell selection.** Starting the SSA with a single cell and simulating the entire population tree is computationally difficult or impossible. We, therefore, track a single cell lineage through time, which allows us to incorporate cell growth and division in stochastic simulation algorithms in an efficient manner. While simulating a single lineage was also done by^41^, we are also able to get the statistical properties of either the whole tree or a sample of extant cells from a single lineage over time. This process is explained in the following section.

**Table S2.** Notations of used variables. For balanced, exponentially-growing cells the volume specific growth rate (U) is equal to the specific growth rate of the population (k).

| Variable | values | pdf | description |
| --- | --- | --- | --- |
| $T$ | $t$ | $\tau(t)$ | Interdivision time |
| $R$ | $r$ | $\rho(r)$ | Partitioning ratio |
| $V$ | | | Cell volume |
| $V_B$ | $v_B$ | $\Psi_B(v_B)$ | Birth volume |
| $V_D$ | $v_D$ | $\Psi_D(v_D)$ | Division volume |
| $V_E$ | $v_E$ | $\Psi_E(v_E)$ | Extant volume |
| $L$ | | | Cell length |
| $L_B$ | $l_B$ | $\ell_B(l_B)$ | Birth length |
| $L_D$ | $l_D$ | $\ell_D(l_D)$ | Division length |
| $L_E$ | $l_E$ | $\ell_E(l_E)$ | Extant length |
| $A$ | $a$ | $u(a)$ | Cell age |
| $k$ | | | Specific growth rate of the population |
| $\mu$ | | | Cell volume growth rate |

#### 7.1 Stochastic simulation of single-cell growth and gene expression matches theory

Within a population of balanced growing cells, three types of samples can be distinguished: samples of extant, mother, and baby cells; their definitions can be found in^27^ and in the main text. The statistics of cell age, generation time and cell volume of these samples are interrelated, for the generation time the following relation holds:

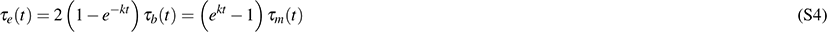

where *k* denotes the specific growth rate of the population and *τ_s_(t*) gives the probability density function of the generation times for the different types of samples (with *s* as either *e*: extant, *b*: baby, or *m*: mother). During lineage simulations we select at each division the daughter cell which will be followed in such a way that the statistics of the resulting lineage corresponds to a sample of mother cells.

To generate a lineage that is representative for a sample of mother cells, at each division the daughter to be followed by the simulation is chosen with a probability according to the fraction of descendants it can be expected to contribute to the population: If daughter 1 is expected to have *n*_1_ descendants at a later time point *t_x_* and daughter two *n*_2_ descendants, the probability *p* to choose daughter 1 is given by

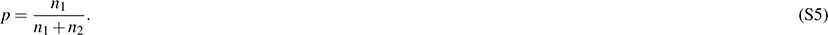

Let *T*_1_ and *T*_2_ be the generation times of daughters 1 and 2, respectively. In balanced growth the number of expected descendants at time *t_x_* is then given by *n*_1_ = *ek*.^(*tx*-*T1*)^ and *n*_2_ = *ek*.^(*tx*-*T2*)^. This is possible because the growth law does not depend on molecule concentrations or previous history. Inserting these relationships for *n*_1_ and *n*_2_ in Eq. (S5) gives

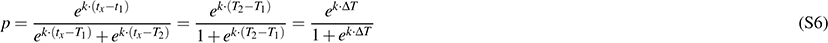

where Δ*T* = *T*_2_ – *T*_1_. In short, the larger daughter cell is more probable to be chosen, because this cell will reach the next division volume sooner and is therefore likely to have more descendants in the population at a later time point.

With as simulation result a lineage that represents a sample of mother cells, statistical properties of other defined samples can be calculated based on the known relationships of generation time and cell age between these samples^27^.

#### 7.2 Conditions to interrelate a lineage and full tree simulation

Here we state the conditions for the simulation of a single lineage that can be used to retrieve the statistics of the entire lineage tree:

1. **The volume at which cells divide has to be independent of the volume at birth, and any other cell-specific properties like concentrations of certain molecules etc.** To ensure that volumes at division are independent of volumes at birth, the distribution of division ratios (ρ) and the distribution of volumes at division (Ψ_*D*_) are chosen such that the largest possible volume at birth is smaller than the smallest possible volume at division (i.e. there is no overlaρbetween the two volume distributions). By default we use beta distributions for both the division ratio as well as the volume at division. Beta distributions are bounded and symmetric if its two positive shape parameters are identical.
2. **The growth law for a single cell has to be deterministic and independent of intracellular concentrations.** The current implementation allows exponential and linear growth laws for single cells. For an exponential growth law the volume specific growth rate (μ) is equal to the specific growth rate of the population (*k*). Otherwise, the specific growth rate of the population needs to be calculated. We do this by using the known relations between the volume at division, volume at birth, the growth law, the volume distribution of extant cells (Equation (S7)^27^), and the fact that this volume distribution has to integrate to one:

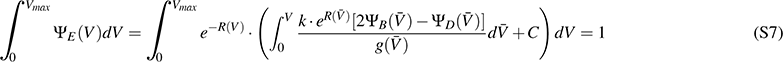

where

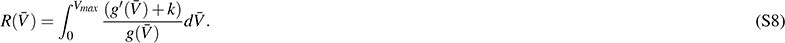

Here *V* is the cell volume, *C* an integration constant (which can be calculated from boundary conditions of the volume distribution), *λ_e_(V)* the volume distribution of a sample of extant cells, Ψ_*B*_(*V*) the volume distribution at birth for a sample of baby cells, Ψ_*D*_(*V*) the volume distribution of a sample of mother cells, and *g'(V)* the differential of the formula of cell volume growth (*g*(*V*) = μ.V for exponential growth and *g*(*V*) = μ for linear volume growth, with μ as the volume specific growth rate). We know Ψ_*D*_(*V*), *g*(*V*), *g*'(*V*), and *C* from the model parameters. Ψ_*B*_(*V*) can be calculated from Ψ_*D*_(*V*) and the partition distribution *ρ(r)* using^28^:

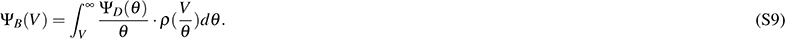 A solution for Eq. (S7) is approximated by using the secant or Newton-Raphson method. For example, we can calculate the copy number distributions for a sample of extant cells (corresponding to any experiment that records molecule copy numbers at a fixed moment in time as for example smFISH) from the simulation of a single lineage by using

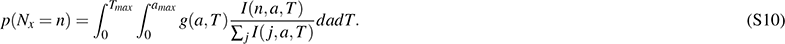

Here, 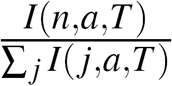 is the relative frequency of copy number equal to *N_x_* in the simulated lineage at age *a* for a cell with generation time *T* approximating *p*(*N*_*x*_ = *n*|*a*, *T*) and *g*(*a*, *T*) is the joint PDF of cell age and generation time,

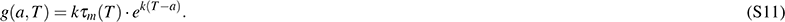

The derivation of *g(a*, *T)* is given in Eq. S13 and S14. Using Eq. (S11) we can rewrite Eq. S10 to:

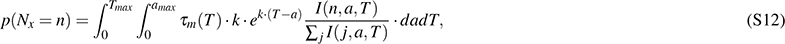

where *I(n*, *a, T)* is an indicator function equal to the number of occurrences of copy number *N_x_ = n* at cell age *a* in a cell with generation time *T*. Obtaining the statistics for an extant cell population consists of two main steps. First, in order to work with the indicator functions, the simulation time series needs to be binned. The binning needs to be performed at regular intervals for cell age. Secondly, a double integral has to be taken which is a slow procedure for infinitesimal small steps of *a* and *T*. We approximate this double integral by using the Riemann sum (or the 2D trapezoidal rule).

Additionally, the statistics for a sample of extant cell volumes can be calculated from the simulation of a single lineage by re-using Eq. S12 and replacing copy numbers with volume data. Calculating the statistics for sample of extant cell volumes requires one additional step—the continuous volume data must be binned to make the distribution discrete.

#### 7.3 The joint distribution of cell age and generation time

In order to obtain the copy number distributions for a sample of extant cells the joint distribution of cell age and generation time for extant cells is required which can be calculated knowing the growth rate and the generation time distribution. The derivation presented here is analogous to the one for the distribution of extant cell generation times in^27^ (p. 528). With a population size of n_0_ at *t =* 0 the rate of formation of new cells equals 2*k* · *n*_0_ · *e^kt^*. The fraction of these cells with generation time smaller than τ which survive at least until time *t* is given by 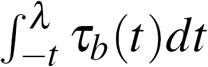 Therefore the total number of cells at time *t* with generation time < λ and age < *a* equals:

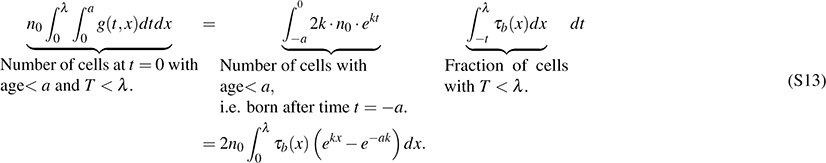

Differentiation yields:

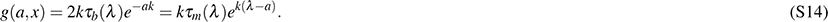

#### 7.4 Illustrations of StochPy simulations with cell growth and division

A typical StochPy modeling session consists of first creating a StochPy cell division model object from a (default) input model. Of course, different user-defined models (in SBML or PySCeS MDL) can be loaded into the model object. Once a model is loaded various simulation parameters that are specific for the SSA with cell growth and division can be set, e.g. μ, Ψ_*D*_, and ρ(see also Tabel S2). Kinetic parameter values and molecule copy numbers can be modified interactively, and simulations can be performed by calling the available analysis methods for model objects. As model objects are fully encapsulated, multiple models can be instantiated from the same (or different) input files at the same time. An example of a short StochPy modeling session within Python is:

~~~
1 import stochpy
2 cmod = stochpy. CellDivision ( model_file = “ImmigrationDeath. psc”)
3 cmod. ChangeParameter(“Ksyn” ,2)
4 cmod. ChangeParameter(“Kdeg” ,0.1)
5 cmod. SetGrowthFunction (growth _r ate = 0.1, grow th_type = “ exponential”)
6 cmod. SetVolumeDistributions (phi = (“beta” ,5 ,5), K= (“fixed” ,0.5) ,
7 phi_beta_mean =2)
8 cmod. DoCellDivisionStochSim (end= 100,mode=“ generations ”)
9 cmod. PlotSpeciesVolumeTimeSeries ()
~~~

where we start by initiating the model object cmod for the immigration-death model depicted in `PySCeS MDL` and modifying the kinetics parameters interactively. Next, we set volume specific growth rate characteristics and volume distributions. Then, we generate one time trajectory with cell growth and division (100 generations) and plot the corresponding (discrete) molecule copy number and volume time series data.

Our lineage corresponds to a sample of mother cells, so we can calculate the statistical properties of e.g. the sample of extant cells via the following high-level functions:

~~~
1 cmod. AnalyzeExtantCells ()
2 cmod. PlotSpeciesExtantDistributions ()
~~~

Here we used the default number of bins for both age and generation times. StochPy generates a warning if the numerical integration was not accurate enough. Choosing a different number of bins for both age and generation time typically solves this issue.

**Table S3.**
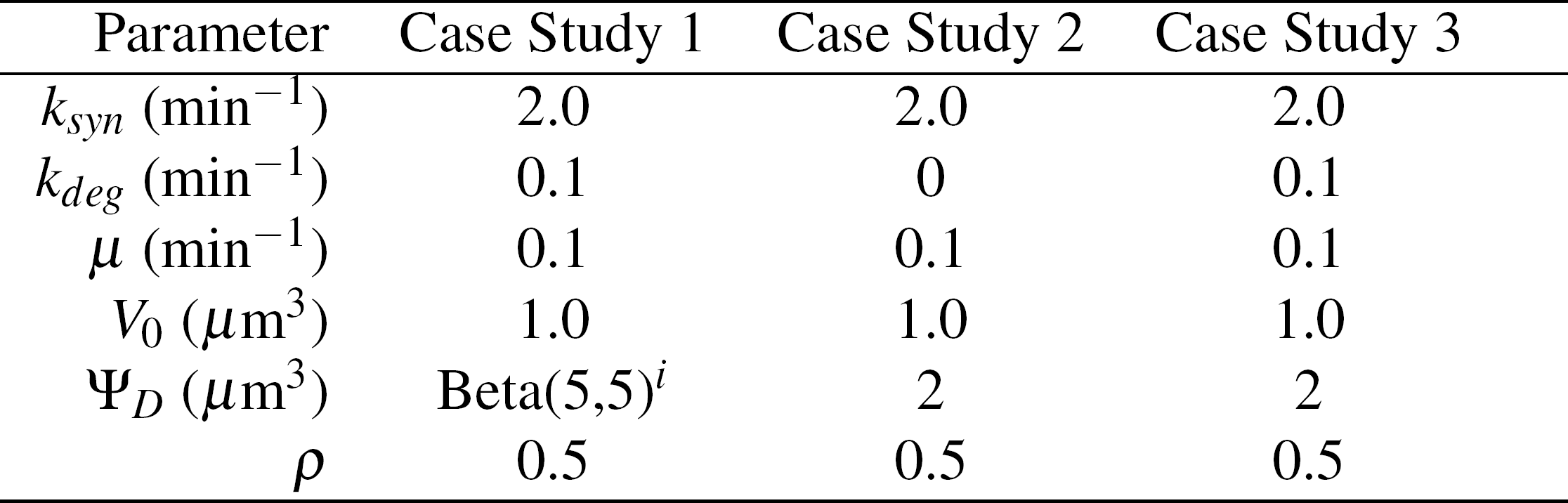
Parameters used in the different case studies. We simulated each SSA with cell growth and division for 10^4^ generations.

| Parameter | Case Study 1 | Case Study 2 | Case Study 3 |
| --- | --- | --- | --- |
| $k_{syn}$ ($\text{min}^{-1}$) | 2.0 | 2.0 | 2.0 |
| $k_{deg}$ ($\text{min}^{-1}$) | 0.1 | 0 | 0.1 |
| $\mu$ ($\text{min}^{-1}$) | 0.1 | 0.1 | 0.1 |
| $V_0$ ($\mu\text{m}^3$) | 1.0 | 1.0 | 1.0 |
| $\Psi_D$ ($\mu\text{m}^3$) | Beta(5,5)<sup>i</sup> | 2 | 2 |
| $\rho$ | 0.5 | 0.5 | 0.5 |

#### 7.5 Comparing StochPy simulations with cell growth and division to analytical solutions

In the following sections, we compare StochPy simulations to theory. In each comparison, we highlight which contributions of cell growth and division to non-genetic variability of cells were included. Besides the simulated results the derived analytical solutions are given. We used the immigration-death model (i.e.“synthesis-degradation”) model to study single-cell transcription with cell growth and division, because this is the only model for which we know the analytical solutions. We used the settings given in Table S3 to—for molecule copy numbers and cell volume—compare time series, distributions at birth *(a =* 0) and division *(a = T*), and distributions for a sample of extant cells.

##### 7.5.1 Case Study 1: Volume statistics are independent of the model

We first analyzed the volume statistics generated by StochPy simulations. While we used the synthesis-degradation model we could have used any model because the volume statistics are completely independent of the model used. Using a different model and/or parameter settings has only an effect on the run time of the simulation. The settings that we used to generate the results are given in the second column of Table S3.

Comparison of the volume statistics generated by StochPy to analytical solutions shows an excellent overall agreement (Fig. S12). In our stochastic simulation we took into account the following contributions of cell growth and division to non-genetic cellular heterogeneity: imprecise, binomial, partitioning of molecules at cell division and heterogeneity in the mother cell volume at cell division. The latter was modeled by drawing division volumes from a Beta distribution (Eq. (S2)). Imprecise volume division from mother to daughter cells was not taken into account because we partitioned the cellular volume exactly between both daughter cells (ρ(0.5) = 1). Heterogeneity in the cell volume at division has several consequences. First, the cellular volume at a given age is variable as is illustrated in Fig. S12A for cells at division *(a = T*) and at birth *(a =* 0). The volume distribution of a sample of extant cells (Fig. S12C) is in agreement with theory. Secondly, the generation time *T* is not deterministic as is illustrated in Fig. S12B and D. In the latter, we compare the generation time distribution of a sample of extant cells (*τ*_e_) obtained by simulation with theory and found an excellent agreement.

**Figure 12.**
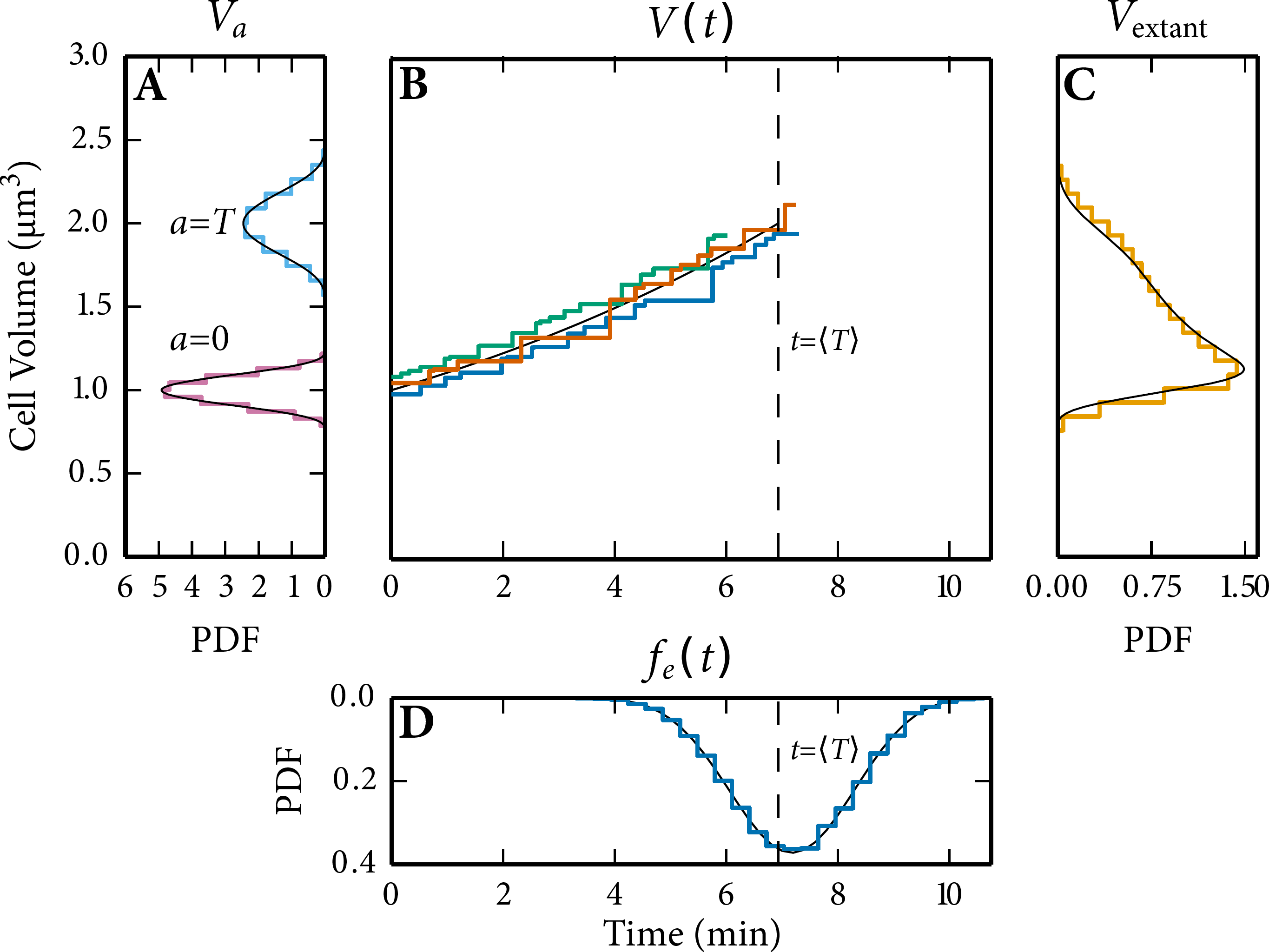
Volume statistics obtained via the SSA with cell growth and division are in agreement with analytical solutions (black). (A) Volume distributions at *a =* 0 and *a = T*. (B) Three stochastic time trajectories of cell volume fluctuate around its analytical solution with V_0_ = 1. Each of the time trajectories has a generation time that is distributed (see panel D) around its mean, 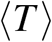 (dashed line). (C) The extant volume distribution. (D) The generation time distribution of a sample of extant cells.

In Fig. S12B we simulated three distinct generations that each started with a different V_0_ drawn from Ψ_*B*_(*V*). We selected the parameter values in our model such that the number of firings (reactions that occur) per generation are limited, such that we can illustrate that StochPy updates *V* only when a reaction fires (Fig. S12B). Our model does not contain any second or higher-order reactions, so this has no effect on the accuracy of the simulation. As explained, this can have an effect if second-order reactions are included and when the rate of firing is slow (then the volume difference becomes significant).

##### 7.5.2 Analytical solutions: Case Study 1 Volume distributions

We drew the cell volume at division (Ψ_D_) from a scaled Beta distribution:

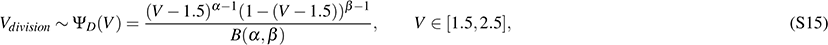

with *V* as the cell volume and *B* as the Beta function that normalizes the distribution. Since the beta distribution is defined on the interval [0,1], we scaled this distribution to the interval [1.5,2.5]. When the division volume was reached, we divided the mature mother cell volume equally between both daughter cells (volume partitioning distribution ρ= 0.5). The distribution of the volume at birth was obtained by calculating the distribution of the product of two random variables, using Ψ_D_ and the partitioning distribution ρ. With ρ as the Dirac delta distribution such that ρ(0.5) = 1. The cell volume distribution at cell birth is then given by:

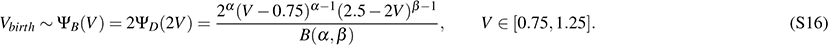

The relationshiρbetween the mean of both distributions is given by

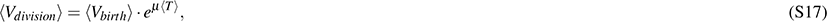

with T = ln(2)/μ thus

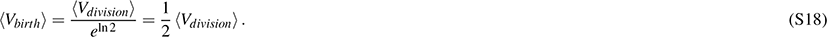

The volume distribution of a sample of extant cells can be calculated as function of Ψ*_b_* and Ψ_*D*_ following the equations deduced by^29^ as shown in^27^:

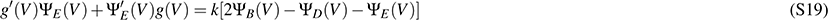

with *g*(*x*) as the growth rate of cells with size *V*, Ψ_*e*_ (*V*) as volume distribution of a sample of extant cells, Ψ_*b*_ as the volume distribution of a sample of baby cells at birth, Ψ_*d*_ as the volume distribution at division of a sample of mother cells, and *k* as the specific growth rate of the population. Assuming *g*(*V*) = *k* . *V*, this function can be simplified to:

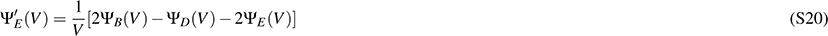

which can be solved to give the volume distribution of a sample of extant cells.

Since *V_division_* and *V_birth_* are independent, the distribution of the ratio ρ= *V_division_/V_birth_* can be calculated from^48^

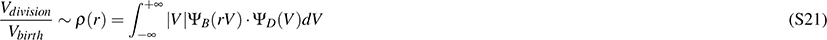

By using the change of variable technique we can transform ρ*(r)* and obtain the distribution of the generation time:

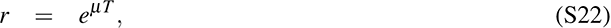

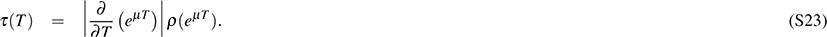

##### 7.5.3 Case Study 2: mRNA synthesis

In addition to predicting accurate volume statistics, we also aimed at predicting accurate molecule copy number statistics when we include the stochastic contributions of cell growth and division to non-genetic cell variability. Unfortunately, we do not know (yet) the analytical solutions if we include all sources of stochasticity of cell growth and division. Hence, we used a deterministic *T* and simplified the model by consideration only first-order synthesis of a molecule and no degradation *(K_deg_ =* 0). A deterministic *T* was achieved by using a fixed Ψ_d_ and ρ*(r)* – the specific settings are given in the third column of Table S3. This means that the only contribution of cell growth and division to non-genetic cell variability that we took into account was the imprecise partitioning of molecules at cell division. The analytical solution for this scenario can be found below.

The results given in Fig. S13 show that predictions made with stochastic simulation are consistent with analytical solutions. More specifically, the mRNA copy number distributions at birth and division obtained with stochastic simulations overlaρwith the theoretical distributions (Fig. S13A) and the time series output of stochastic simulations fluctuate, as expected, around the theoretical time series (Figu. S13B). Here, the deviation is explained by inherent stochasticity in net molecule synthesis in stochastic simulations. We also found an excellent agreement between the simulated and theoretical molecule copy number distribution of a sample of extant cells (Fig. S13C). This result demonstrates that we can simulate a single lineage that represents a sample of mother cells and get information about other samples such as the molecule copy number distribution of a sample of extant cells.

##### 7.5.4 Analytical solutions: Case Study 2 mRNA synthesis

In the second case study, we used a zero-order mRNA synthesis model in exponentially growing cells. With a deterministic generation time, the theoretical mRNA copy number distributions of this model are known^22^.

###### Average molecule copy numbers at birth and division

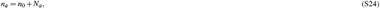

The mRNA copy number in a cell at a certain age (a) is given by

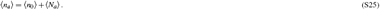

where *n_a_* is the mRNA copy number at cell age *a, n_0_* as the mRNA copy number obtained at cell division, and *N_a_* as the mRNA molecules synthesized since the last cell division at age *a.* The following relationshiρholds for the averages

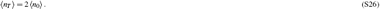

When symmetric partitioning at the end of a cell cycle (t = *T*) is assumed, the average mRNA copy number at birth is half the average of mRNA copy number at division:

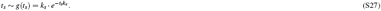

This means that cells double their average number of molecules during one cell cycle *(a =* 0 to *a = T*). The average amount of molecules produced at *t = T* equals the average number molecules obtained at *t =* 0.

**Figure S13.**
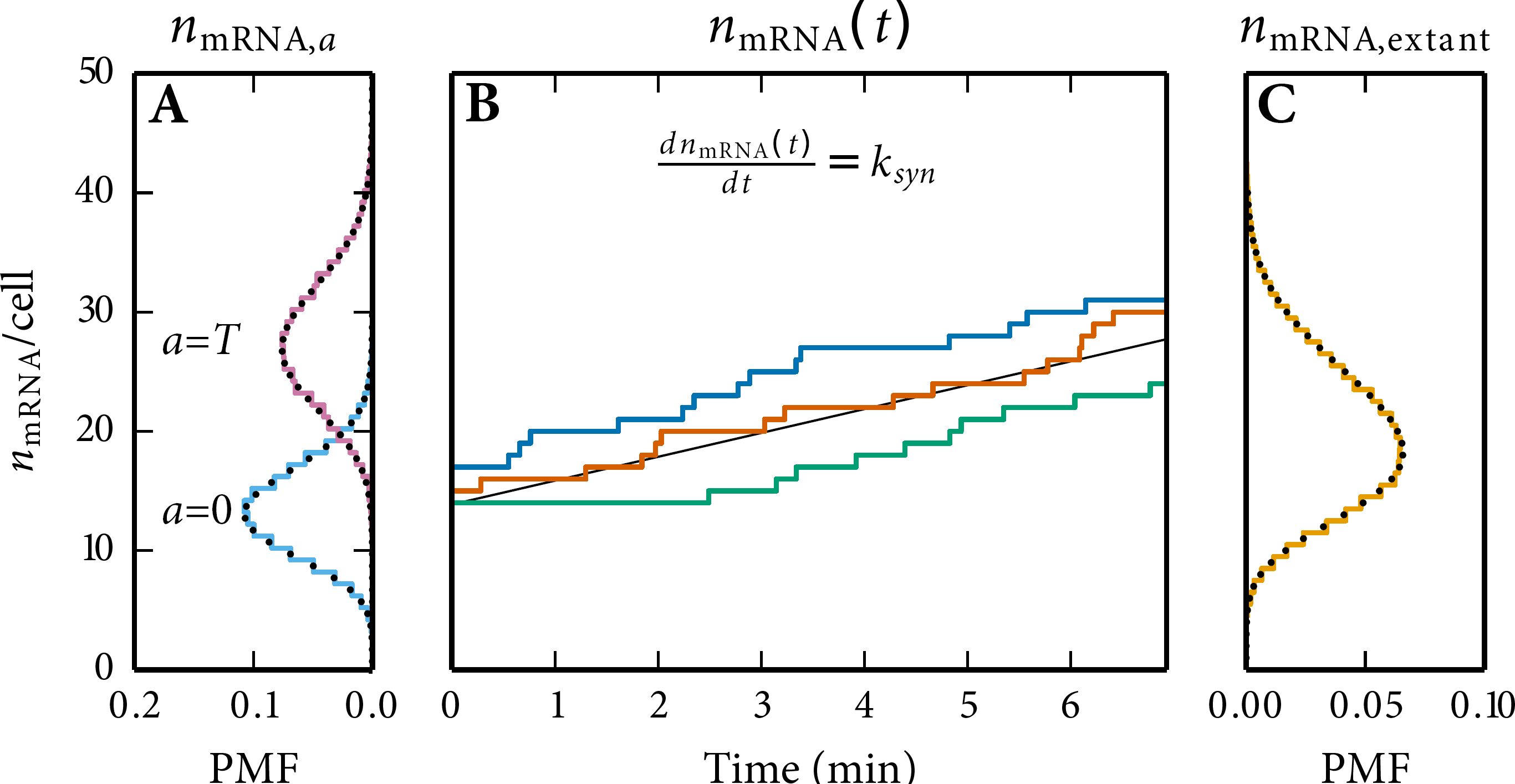
Molecule copy number statistics obtained for the mRNA synthesis model, via the SSA with cell growth and division, is in agreement with analytical solutions. Simulation results are shown in colour, and analytical solutions in black. (A) *n_mRNA_* copy number distributions at *a =* 0 and *a = T*. (B) three stochastic time trajectories of n_mRNA_. (C) the extant *n_mRNA_* copy number distribution.

##### 7.5.5 Poisson distributed molecule copy numbers at a specific age

The waiting time distribution of a first-order synthesis process is exponentially distributed, the time between consecutive events is:

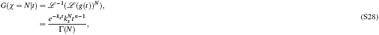

The time to make *N* molecules follows a gamma distribution and can be derived from the N-th convolution of *g(t)* using the generating function, which is the Laplace transform (*L*):

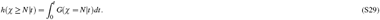

with Г [.] as the incomplete gamma function. The probability to produce more than *N* molecules in time *t* equals:
The probability mass function for the production of *N* molecules at time *t* is given by:

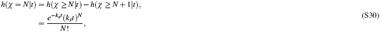

which is a Poisson distribution with mean *k_s_t*. Of course, time can be interchanged by age, *t = a.*

##### 7.5.6 mRNA copy number distribution of a sample of extant cells

The molecule copy number distribution of a sample of extant cells was earlier determined by^22^

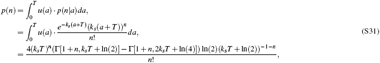

with Г[.] as the incomplete gamma function, *u(a)* as the cell age distribution for a population of cells with a deterministic generation time *T*, and *k*_*s*_ as the synthesis rate constant.

##### 7.5.7 Case Study 3: mRNA synthesis and degradation

Proteins are actively degraded into the cell, so we decided to extent our simple model with active degradation. Except for the degradation rate, we used exactly the same settings (fourth column of Table S3) as in the example without active degradation. The imprecise partitioning of molecules at cell division was, therefore, again the only contribution of cell growth and division to non-genetic variability that was taken into account.

The results given in Fig. S14 show again an almost perfect agreement between stochastic simulation and analytical solutions (species distribution of a sample of extant cells was numerically solved). The corresponding analytical solution can be found below. Comparison of Figs. S13 and S14 shows that, as expected, adding active degradation results in lower mRNA copy numbers at different ages in the lineage and also in the extant cell population.

**Figure S14.**
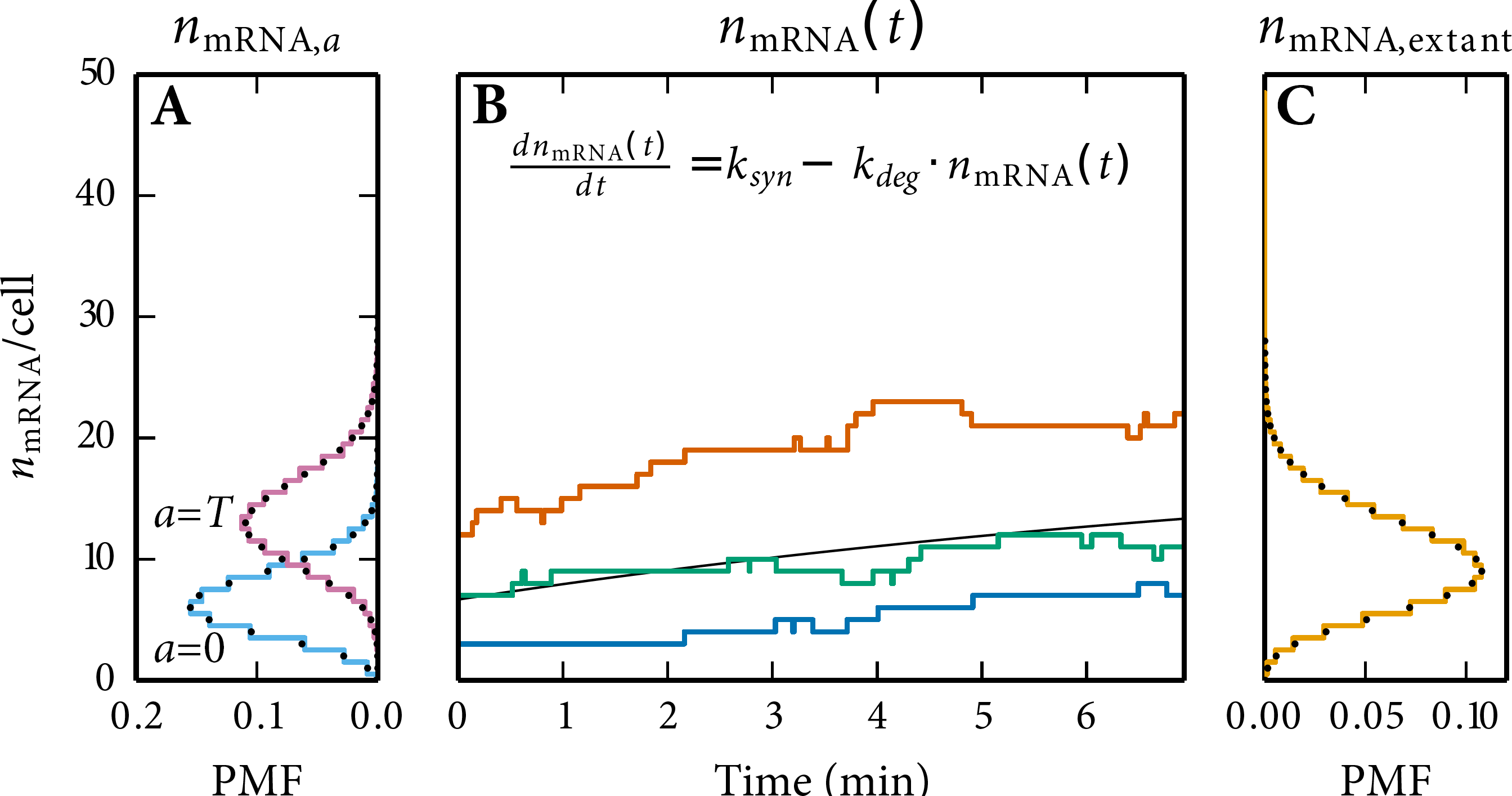
Molecule copy number statistics obtained for the mRNA synthesis and degradation model via the SSA with cell growth and division are in agreement with analytical solutions. Simulation results are shown in colour, and analytical solutions in black. (A) *n_mRNA_* copy number distributions at *a =* 0 and *a = T*. (B) three stochastic time trajectories of *n_mRNA_* fluctuate around its analytical solution. (C) the extant *n_mRNA_* copy number distribution.

##### 7.5.8 Analytical solutions: Case Study 3 mRNA synthesis and degradation

In the third case study, we considered a model consisting of a zero-order mRNA synthesis reaction and first order mRNA degradation reaction, in exponentially growing cells. When we assume a deterministic generation time, the theoretical molecule copy number distributions at a given age of this model are known^22^.

##### 7.5.9 Poisson distributed molecule copy numbers at a specific age

With a synthesis and degradation reaction, the average mRNA copy number in a cell of a given age *a* depends on the mRNA copy number at *t =* 0 and the *net* synthesis until time a. For a linear model with an constant production and degradation rate the average copy number can be written as

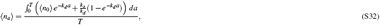

where *T* is the (deterministic) generation time (T = ln(2)/u) and (n_0_) as:

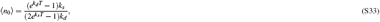

The copy number distribution at age *a* is given by a Poisson distribution (see^22^ for derivation)

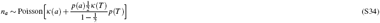

with the number of molecules produced during a cell cycle as uρuntil age *a* as 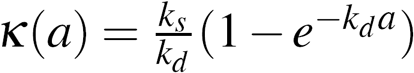 and the survival probability of molecules as *p(a) = e^k_d_a^.*

##### 7.5.10 Molecule copy-number distribution of a sample of extant cells

The molecule copy-number distribution of the sample of extant cells was determined by numerical solving:

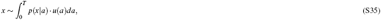

with the *p*(*x*|*a*) determined by Eq. (S34) and the age distribution, *u*(*a*), taken from^27^ in combination with a deterministic generation time, which then for *u(a)* gives:

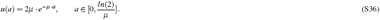

